# CCAFE: Estimating Case and Control Allele Frequencies from GWAS Summary Statistics

**DOI:** 10.1101/2024.10.24.619530

**Authors:** Hayley R Stoneman, Adelle Price, Christopher R Gignoux, Audrey E Hendricks

## Abstract

Methods involving summary statistics in genetics can be quite powerful but can be limited in utility. For instance, many post-hoc analyses of disease studies require case and control allele frequencies (AFs), which are not always published. We present two frameworks to derive case and control AFs from GWAS summary statistics using the odds ratio, case and control sample sizes, and either the total (case and control aggregated) AF or standard error (SE). In simulations and real data, derivations of case and controls AFs using total AF is highly accurate across all settings (e.g., minor AF, condition prevalence). Conversely, derivations using SE underestimate common variant AFs (e.g. minor allele frequency >0.3) in the presence of covariates. We develop an adjustment using gnomAD AFs as a proxy for true AFs, which reduces the bias when using SE. While estimating case and control AFs using the total AF is preferred due to its high accuracy, estimating from the SE can be used more broadly since SE can be derived from p-values and beta estimates, which are commonly provided. The methods provided here expand the utility of publicly available genetic summary statistics and promote the reusability of genomic data. The R package *CCAFE*, with implementations of both methods, is freely available on Bioconductor and GitHub.

## 1. Introduction

Growth in the field of genetic and genomic research has rapidly increased the quantity of genome-wide association studies (GWAS) as well as the availability of datasets from these studies. These data are often made publicly available through summary statistics to ease storage and privacy concerns. This increase in summary-level data has catalyzed the development of new analyses and methods such as GWAS meta-analysis (Tam et al., 2019), polygenic risk scores (PRS) (Kullo et al., 2022; van Rheenen et al., 2019), Mendelian Randomization (MR) (Sanderson et al., 2022), and external common controls (Hendricks et al., 2018; Lee et al., 2017; Wojcik et al., 2022). GWAS summary statistics often include a subset of odds ratio (OR) or effect size (beta), standard error (SE), p-value, allele frequency (AF), and sample size. However, there is a lack of consistency and standardization for reporting summary statistics, in both content and format, presenting challenges in data reuse (Lyon et al., 2021; Matushyn et al., 2022; Thelwall et al., 2020). A recent study of 327 summary statistics files found over 100 unique formats (Murphy et al., 2021) resulting from different types of traits studied (i.e. binary or quantitative), the software used for analysis, or simply author choice. While movements have been made to standardize summary statistics reporting (Buniello et al., 2019; Lyon et al., 2021), inconsistency in the reporting can limit, and sometimes fully hinder, the use of these data for further studies. In an assessment of the 2021 requirements for submission to the GWAS catalog, a recent study found that over 50% of studies submitted between January 2020 and July 2022 were missing at least one mandatory field (Hayhurst et al., 2023). Notably, the most commonly missing fields were effect AF (>40% missing) and SE (>25% missing)- although SE can be recapitulated from the effect estimate and p-value. Inclusion of case and control AFs was even more rare. Implementation and enforcement of these standards has the potential to double the number of usable datasets for downstream analyses.

Further, clear standards for reporting the effect AF do not exist, with researchers frequently summarizing across all individuals to report the whole sample AF from GWAS, resulting in the aggregation of case and control samples. Without access to case and control specific AFs, secondary uses for summary data like group PRS (Yang et al., 2022) and as external controls may not be possible. In 2022, Yang et al. presented a framework to reconstruct GWAS case and control AFs using the GWAS case and control sample sizes, OR, and SE as part of a GWAS meta-analysis software called ReACt. While Yang et al. evaluated the use of their derived case and control AF in secondary analyses such as case-case GWAS and meta-analysis, there was no direct evaluation of the derived case-control AFs.

Here, we develop a method to estimate case and control AF using OR, case and control sample size, and total AF. In real data and simulations, we evaluate this method as well as Yang et al.’s case and control AF derivation that uses SE with and without a bias adjustment. The original framework using SE was published by Yang *et al*. as part of the ReACt software. However, ReACt does not have an independent function to calculate the case and control AFs. Therefore, we provide methods in the *CCAFE R* software package (R Core Team, 2024), available on GitHub and Bioconductor (Huber et al., 2015). *CCAFE* enables derivation of unbiased GWAS case and control AFs using the GWAS total sample AF or SE, number of cases and controls, and effect estimate, supporting broader use of GWAS summary statistics. These well-documented and user-friendly functions will help to expand the use of summary statistics.

## 2. Methods

### 2.1 Implementation

#### 2.1.1 CaseControl_AF mathematical framework

We derive case and control AFs for a given variant *i* using the case and control sample sizes (*N*_*case*_ and *Nc*_*control*_), *OR*_*i*_, and total AF (*AF*_*total,i*_). *AF*_*total,i*_ and *OR*_*i*_ can be represented as shown in equations (1) and (2).

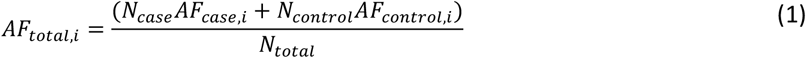

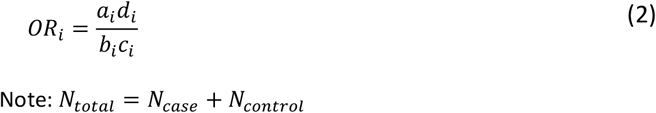

Here *a, b, c*, and *d* are the cells of a two-by-two contingency table representing the allele counts of the effect and non-effect, or alternate and reference, alleles for cases and controls. These quantities can be calculated as shown in equations (3) – (6).

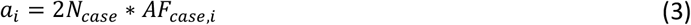

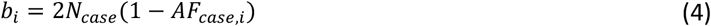

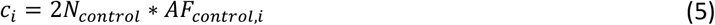

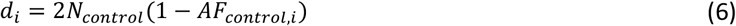

Substituting equations (3) through (6) into the OR equation in (2), we arrive at equation (7).

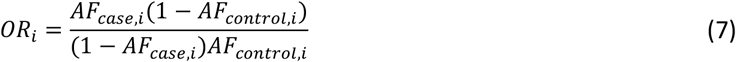

We can then use equation (1) to solve for *AF*_*case,i*_ (shown in equation (8)) and substitute it into equation (7), which results in a quadratic as shown in equation (9).

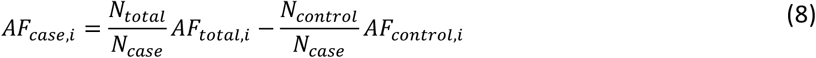

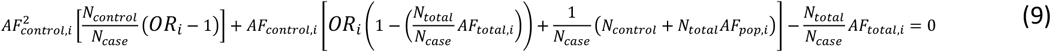

We solve for the quadratic roots and choose *AF*_*control,i*_ to be the root between 0 and 1. The full derivation is shown in the supplement. Additionally, we show there is only one solution for *AF*_*control,i*_ that falls between the bounds of 0 and 1 (**Figure S1, Table S1**). *AF*_*case,i*_ is then estimated using equation (8).

#### 2.1.2 CaseControl_SE mathematical framework

The full derivation for the case and control AFs using SE can be found in the supplemental information of the original publication by Yang *et al*. Briefly, the method relies on allele counts *a, b, c*, and *d* as shown in equations (3) – (6). As this results in four unknown quantities, four equations are used to solve the system. These four unknown quantities are related to SE, the sample size of cases and controls, and OR. Ultimately, Yang et al. solve for *d* and use the resulting quadratic to solve for the allele counts (*a, b, c, d*) and then the AFs. To fulfill the assumption that the solution is between [0, 1], Yang et al. use the minor allele frequency (MAF) resulting in a loss of connection with an allele. While the minor and major alleles can likely be accurately inferred when the MAF is very different from 0.5, as the MAF approaches 0.5 accurate inference of the minor allele becomes difficult. Notably, the resulting quadratic solved in this framework utilizes the total allele number (i.e. 2*N for autosomes) in cases and controls, rather than the allele frequency. This results in the need for sex chromosome-specific implementations, and thus requires information regarding the number of X and Y chromosomes in the case and control samples.

#### 2.1.3 CaseControl_SE bias correction

To enable unbiased estimation of case and control AFs using SE, we present a method to estimate and adjust for the bias. We use gnomAD v3.1.2 MAFs as proxies for the true total sample MAFs and fit a second order polynomial regression for five MAF bins ([0.0, 0.1) [0.1, 0.2) [0.2, 0.3) [0.3, 0.4) [0.4, 0.5]). The estimated MAF is the outcome, and the proxy MAFs from gnomAD are the predictors. Comparing this predicted MAF estimate value to the proxy MAF, we estimate the bias across the MAF spectrum. The fitted model is used to remove bias from all GWAS variant case and control AF estimates, even those not represented in gnomAD (**Figure S2**). Full details of the bias correction framework are available in the supplement.

#### 2.1.4 Implementation in R

We include the method from Yang *et al*. as a function, named CaseControl_SE, in our R package. We also include our framework proposed here as the function CaseControl_AF. Both methods (using either total AF or SE) rely on solving the roots of a quadratic. As we know that these quantities (AFs) exist and are real numbers, this can be done easily in R using the quadratic formula and is scalable for large datasets. Both methods use the number of cases and controls, ORs, and either total AF or SE as input. Note, the OR can be derived from the beta estimate and the SE can be derived from the effect estimate and either the p-value or the test statistic. To use the SE to derive the MAF for sex chromosomes, the user must also include the number of X and Y chromosomes per case and control sample.

Additionally, the user may obtain corrected MAF estimates from CaseControl_SE by including a data frame containing variants that have been harmonized between the observed data and a proxy dataset such as gnomAD. This data frame must contain variant information (chromosome and position) as well as the proxy MAF. This data frame does not need to contain all variants in the observed dataset. The adjusted MAFs are appended as three additional columns to the original data input.

### 2.2 Simulation study

We used the R package *PhenotypeSimulator* (Meyer & Birney, 2018) to simulate models with genotypes for 10,000 variants (of which 100 were causal), one binary phenotype, and zero, one, or three covariates (**Figure S3**). *PhenotypeSimulator* generates phenotypes as the sum of genetic variant effects, covariate effects, and observational noise. We simulated multiple sample sizes (n=1000, 6000, 10000, 50000, 100000) with equal cases and controls as well as imbalanced sample sizes (600 cases and 5400 controls to emulate the Pan-UK Biobank African Diabetes sample, and 1200 cases and 48800 controls). When including one covariate, the covariate was simulated to be similar to biological sex as a binary variable with probability of 0.5. For three covariates, two additional variables were simulated, one categorical variable with five categories and one normally distributed with mean of 30 and standard deviation of 20. The binary phenotype was simulated given genotypes, covariates, and random noise. Then the binary phenotype was used to define cases, where the phenotype was coded as ‘1’, and controls, where the phenotype was coded as ‘0’. The definition of case and control status by binary phenotype was used to split the simulated genotypes into cases and controls, from which case and control AFs were calculated. The genotype, phenotype, and covariate data were used to fit a logistic regression model from which the OR and SE for each simulated genetic variant were obtained for further use in estimating case and control AF (additional details in data availability).

The summary statistics from logistic regressions (e.g., OR and SE) of the simulated data were used in *CCAFE* to estimate the case and control AF. The estimated AFs were compared to the simulated AFs and accuracy and variability were assessed using Lin’s Concordance Correlation Coefficient (Lin, 1989).

### 2.2 Real data application

#### 2.2.1 Publicly available datasets

We tested the ability of the methods to reconstruct case and control AFs from real datasets for which case and control AFs were made publicly available (**Table 1**). These datasets had a range of case and control sample sizes, number of genetic variants, and covariates included in the original GWAS. We used 148 variants from a 2018 prostate cancer PGS, which was generated from the data collected for a GWAS of 79,148 cases and 61,106 controls (Schumacher et al., 2018). This GWAS published control AFs, which were used with the OR to derive the case AFs (details in the supplement). Additionally, we use the Pan-UK Biobank (Pan-UKB) GWAS summary statistics for diabetes in European (EUR) and African (AFR) samples (Pan-UKB team, 2020). Here 9,178,564 genome-wide variants were available. The Pan-UKB EUR GWAS contained 16,550 cases and 403,923 controls, while Pan-UKB AFR GWAS contained 668 cases and 5956 controls.

**Table 1.** Lin’s Concordance Correlation Coefficient for the two methods included in *CCAFE* (CaseControl_AF and CaseControl_SE) in simulations.

|  |  |  |  | Lin's CCC |  |  |  |
| --- | --- | --- | --- | --- | --- | --- | --- |
| Cases | Controls | Total | Covariates | Cases |  | Controls |  |
|  |  |  |  | AF | SE | AF | SE |
| 500 | 500 | 1000 | 0 | 0.9916 | 0.9811 | 0.9917 | 0.9811 |
|  |  |  | 1 | 0.9905 | 0.7430 | 0.9914 | 0.7420 |
|  |  |  | 3 | 0.9916 | 0.6913 | 0.9905 | 0.6899 |
| 3000 | 3000 | 6000 | 0 | 0.9982 | 0.9952 | 0.9983 | 0.9951 |
|  |  |  | 1 | 0.9981 | 0.8457 | 0.9980 | 0.8442 |
|  |  |  | 3 | 0.9981 | 0.6690 | 0.9982 | 0.6690 |
| 5000 | 5000 | 10000 | 0 | 0.9988 | 0.9967 | 0.9988 | 0.9966 |
|  |  |  | 1 | 0.9987 | 0.8607 | 0.9988 | 0.8599 |
|  |  |  | 3 | 0.9986 | 0.6747 | 0.9986 | 0.6747 |
| 25000 | 25000 | 50000 | 0 | 0.9995 | 0.9989 | 0.9995 | 0.9989 |
|  |  |  | 1 | 0.9993 | 0.8641 | 0.9993 | 0.8643 |
|  |  |  | 3 | 0.9994 | 0.6806 | 0.9993 | 0.6808 |
| 50000 | 50000 | 100000 | 0 | 0.9995 | 0.9993 | 0.9995 | 0.9993 |
|  |  |  | 1 | 0.9995 | 0.7589 | 0.9994 | 0.7593 |
|  |  |  | 3 | 0.9995 | 0.6761 | 0.9995 | 0.6754 |
| 600 | 5400 | 6000 | 0 | 0.9921 | 0.9941 | 0.9980 | 0.9882 |
|  |  |  | 1 | 0.9916 | 0.8410 | 0.9980 | 0.8369 |
|  |  |  | 3 | 0.9918 | 0.8234 | 0.9980 | 0.8189 |
| 1200 | 48800 | 50000 | 0 | 0.9956 | 0.9992 | 0.9994 | 0.9955 |
|  |  |  | 1 | 0.9954 | 0.9903 | 0.9994 | 0.9868 |
|  |  |  | 3 | 0.9954 | 0.9773 | 0.9994 | 0.9731 |

All simulations contained 10,000 variants (100 causal). AF and SE columns contain Lin's CCC between estimated and true case and control AF for CaseControl\_AF and CaseControl\_SE, respectively

#### 2.2.2 Correction using gnomAD as proxies

We applied our bias correction to the genome-wide Pan-UKBB diabetes GWAS data (>9 million variants), using chromosome one variants from gnomAD as proxies. We lifted over Pan-UKBB chromosome one positions from GRCh37 to GRCh38 using LiftOver (Hinrichs et al., 2006) and merged the lifted over Pan-UKBB data with gnomAD v3.1.2 (Karczewski et al., 2020) retaining the intersection of variants contained in both datasets. We removed variants where the gnomAD variant alleles did not match either allele in Pan-UKBB and flipped the alleles of variants for which allele frequency was calculated on opposing alleles. The final dataset contained 1,212,618 variants on chromosome one. We used the gnomAD non-Finnish European (NFE) as the proxy total MAFs for the Pan-UKBB EUR sample, and the gnomAD African/African American (AFR/AFRAM) as the proxy MAFs for the Pan-UKBB AFR sample. We then adjusted the estimated MAFs for all >9 million genome-wide variants in each sample.

## 3 Results

### 3.1 Simulation Results

For both methods (i.e., using total AF or SE) we found that the variability of the estimates compared to the true MAF increases with MAF and decreases as sample size increases (**Figure 1, Figure S4-S7**). When covariates are included in the regression, CaseControl_SE has increasing bias (average difference between the estimated and true MAF) as the MAF increases with MAF being systematically underestimated. Conversely, CaseControl_AF provides accurate estimates across the MAF spectrum in all tested scenarios even with the presence of covariates. This is further shown by the Lin’s CCC between the true MAF and estimated MAF, which decreases as the number of covariates increases for CaseControl_SE but remains near 1 for CaseControl_AF (**Table 1**). Notably, for both methods, the causal variant case and control AF estimates are more variable compared to the estimates for the non-causal variants (**Figure 1, Figure S4-S7**). We also found that using total MAF rather than total AF added additional variability in the estimates from CaseControl_AF (**Figure S8**).

**Figure 1.**
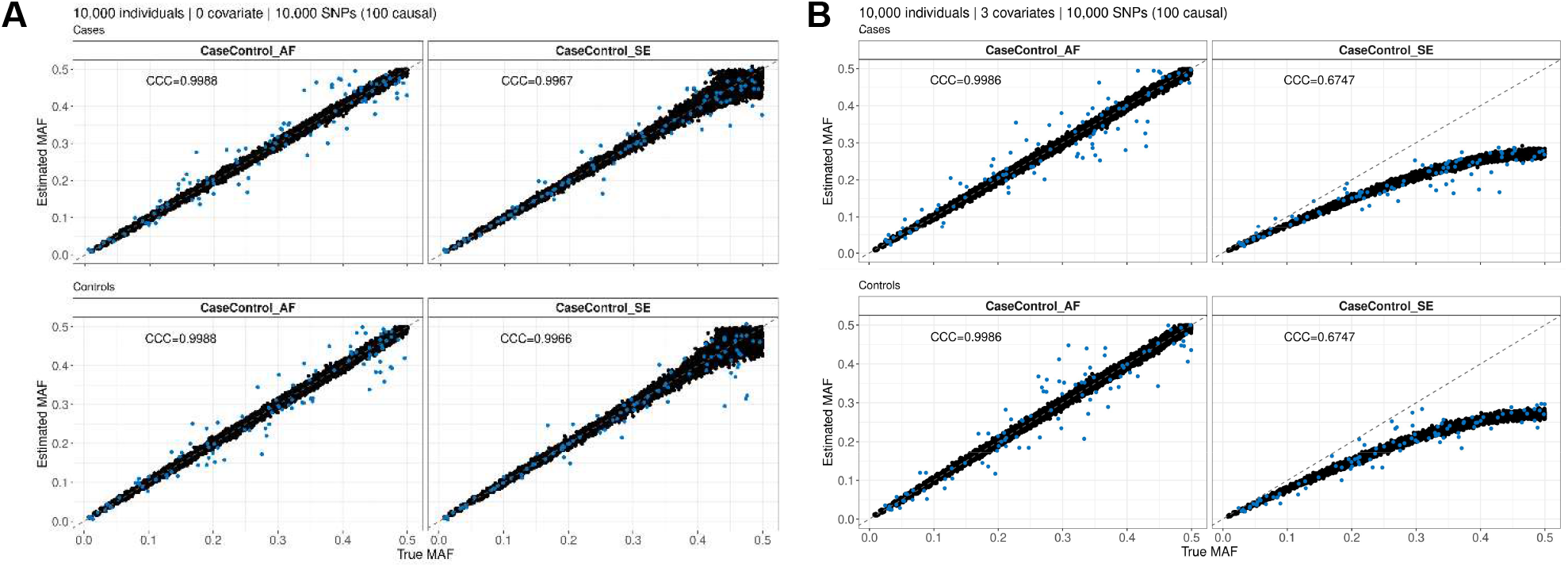
Estimated case and control AFs from summary statistics. Simulated genotypes and phenotypes were generated using the PhenotypeSimulator R package. Genotypes for 10,000 variants, of which 100 were causal (shown in blue), were generated for 5,000 cases and 5,000 controls. Logistic regression was used along with 0 (**A**) or 3 (**B**) covariates to generate per variant summary statistics. CaseControl_AF and CaseControl_SE methods were used to estimate the case and control AFs. Using CaseControl_SE, bias was observed at higher MAFs when covariates were included with a systematic underestimation of MAFs (**B**). CaseControl_AF was accurate across the MAF spectrum, regardless of whether covariates were included. Lin’s CCC is shown between the true simulated MAF and the estimated MAF.

### 3.2 Real data analysis

CaseControl_SE underestimated the true case and control AFs in all datasets, especially for variants with a higher MAF (**Figure 2A**). We identified the highest bias for variants with small SE, in both absolute and relative terms (**Figure S9-S10**). Variants with smaller SE had higher bias, which were often those with a large MAF (e.g., MAF > 0.3). Additionally, across studies, the minimum SE was smaller when case sample sizes were larger, resulting in higher bias for CaseControl_SE in case and control AFs estimated in datasets with larger case sample sizes (**Figure S9-S10**). When using SE to reconstruct the case and control AF, the CCC between the true MAF and estimated MAF decreased as MAF increased in all datasets and was compounded for datasets with larger numbers of cases (**Table 2-3**). Conversely, CaseControl_AF accurately reconstructed the case and control AFs in all datasets (**Figure 2B**) and the CCC of the estimated case and control AF with the true AFs was at or near 1 (**Table 2-3**).

**Table 2.** Lin’s Concordance Correlation Coefficient for CaseControl_AF, CaseControl_SE, and bias corrected CaseControl_SE between true and estimated case MAF.

|  |  |  |  |  | Lin's CCC |  |  |
| --- | --- | --- | --- | --- | --- | --- | --- |
| Trait | Cases | Controls | Bin | N Variants | AF | SE | SE Corrected |
| Prostate Cancer | 79148 | 61106 | [0.0, 0.1] | 18 | 1 | 0.9643 | NA* |
|  |  |  | (0.1, 0.2] | 27 | 1 | 0.8560 | NA |
|  |  |  | (0.2, 0.3] | 35 | 1 | 0.6592 | NA |
|  |  |  | (0.3, 0.4] | 32 | 1 | 0.2401 | NA |
|  |  |  | (0.4, 0.5] | 36 | 1 | 0.0596 | NA |
| PanUKBB Diabetes | 16550 | 403923 | [0.0, 0.1] | 3721679 | 0.99998 | 0.9701 | 0.9651 |
|  |  |  | (0.1, 0.2] | 1766962 | 0.99993 | 0.7346 | 0.9805 |
|  |  |  | (0.2, 0.3] | 1355742 | 0.99989 | 0.3977 | 0.9404 |
|  |  |  | (0.3, 0.4] | 1203911 | 0.99986 | 0.1425 | 0.8244 |
|  |  |  | (0.4, 0.5] | 1130270 | 0.99985 | 0.0213 | 0.2022 |
| PanUKBB Diabetes | 668 | 5956 | [0.0, 0.1] | 3453818 | 0.99871 | 0.9624 | 0.9774 |
|  |  |  | (0.1, 0.2] | 1871888 | 0.99640 | 0.7179 | 0.8891 |
|  |  |  | (0.2, 0.3] | 1487374 | 0.99500 | 0.4168 | 0.7523 |
|  |  |  | (0.3, 0.4] | 1242683 | 0.99409 | 0.1666 | 0.4580 |
|  |  |  | (0.4, 0.5] | 1126168 | 0.99358 | 0.0278 | 0.2146 |

\* Prostate cancer MAFs were not corrected using gnomAD as proxies due to small number of overlapping variants with gnomAD

**Table 3.** Lin’s Concordance Correlation Coefficient for CaseControl_AF, CaseControl_SE, and bias corrected CaseControl_SE between true and estimated control MAF.

| Trait | Cases | Controls | Bin | N Variants | AF | Lin's CCC |  |
| --- | --- | --- | --- | --- | --- | --- | --- |
|  |  |  |  |  |  | SE | SE Corrected |
| Prostate Cancer | 79148 | 61106 | [0.0, 0.1] | 18 | 1 | 0.90315 | NA* |
|  |  |  | (0.1, 0.2] | 27 | 1 | 0.73818 | NA |
|  |  |  | (0.2, 0.3] | 35 | 1 | 0.50244 | NA |
|  |  |  | (0.3, 0.4] | 32 | 1 | 0.24222 | NA |
|  |  |  | (0.4, 0.5] | 36 | 1 | 0.03586 | NA |
| PanUKBB Diabetes | 16550 | 403923 | [0.0, 0.1] | 3721679 | 1 | 0.9700 | 0.9651 |
|  |  |  | (0.1, 0.2] | 1766962 | 1 | 0.7338 | 0.9802 |
|  |  |  | (0.2, 0.3] | 1355742 | 1 | 0.3972 | 0.9389 |
|  |  |  | (0.3, 0.4] | 1203911 | 1 | 0.1428 | 0.8214 |
|  |  |  | (0.4, 0.5] | 1130270 | 1 | 0.0213 | 0.2060 |
| PanUKBB Diabetes | 668 | 5956 | [0.0, 0.1] | 3453818 | 0.99998 | 0.9567 | 0.9760 |
|  |  |  | (0.1, 0.2] | 1871888 | 0.99995 | 0.7097 | 0.8850 |
|  |  |  | (0.2, 0.3] | 1487374 | 0.99994 | 0.4129 | 0.7449 |
|  |  |  | (0.3, 0.4] | 1242683 | 0.99993 | 0.1651 | 0.4508 |
|  |  |  | (0.4, 0.5] | 1126168 | 0.99992 | 0.0281 | 0.2175 |

\* Prostate cancer MAFs were not corrected using gnomAD as proxies due to small number of overlapping variants with gnomAD

**Figure 2.**
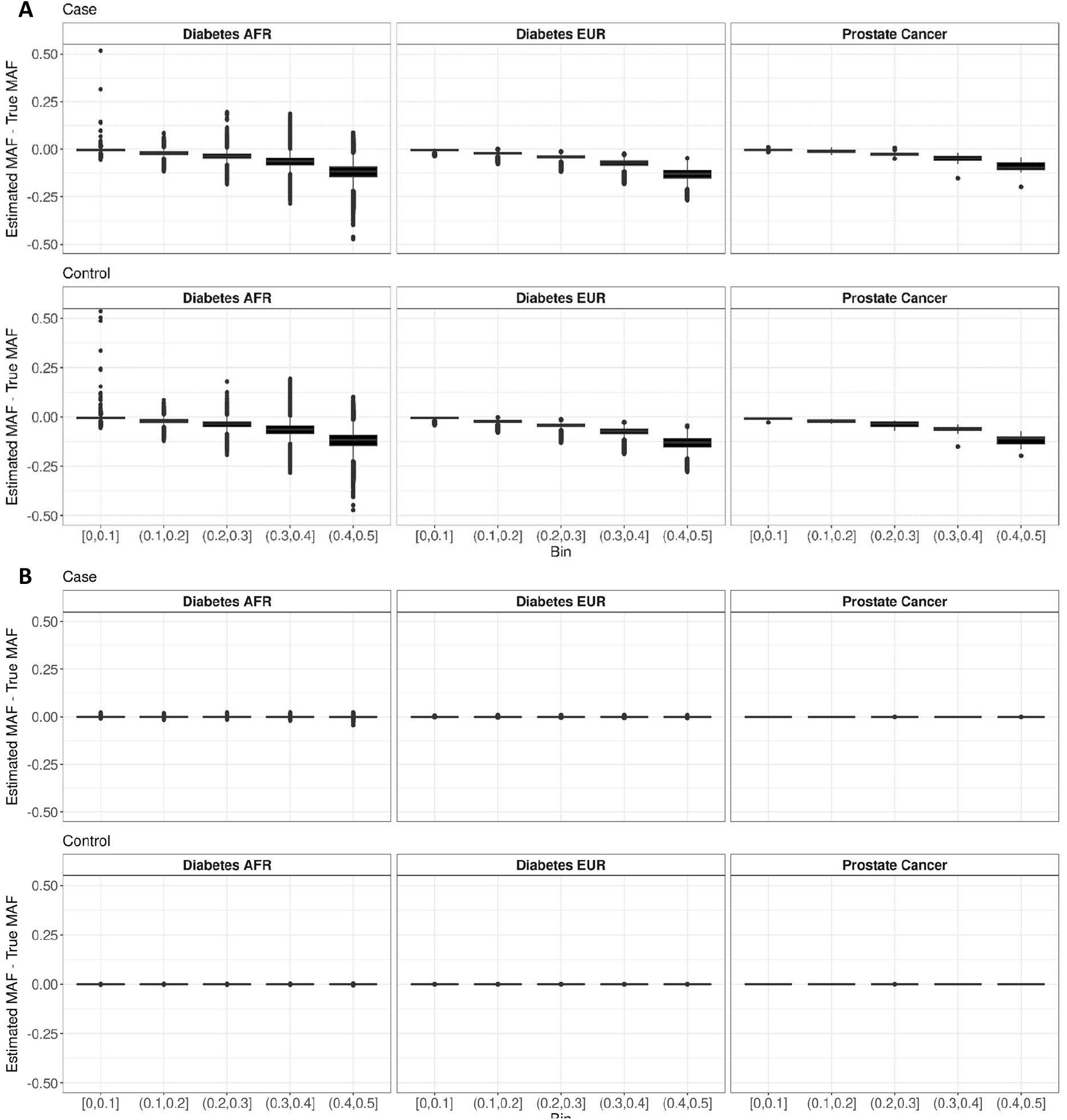
Comparison of case and control AF estimation in multiple real datasets. Results of estimating the case MAFs for six datasets with various sample sizes using (**A**) CaseControl_SE, the method proposed in the ReACt software using SE, and (**B**) CaseControl_AF, the framework developed here using total AF. The Prostate Cancer dataset (Ncase=79148; Ncontrol=61106) has 148 variants from a 2018 PRS, and the true case and control AFs were published as part of the discovery GWAS. Diabetes EUR (Ncase=16550; Ncontrol=403923) and Diabetes AFR (Ncase=668; Ncontrol=5956) contain >9 million variants from the PanUKBB GWAS. CaseControl_SE underestimates the true MAF, with bias increasing and precision (width of the boxplot) decreasing as the true MAF increases. Conversely, we see highly accurate estimation of known AFs using CaseControl_AF, with some variability in datasets with small sample sizes.

### 3.3 CaseControl_SE bias correction

After observing the bias between the true and estimated MAF for the CaseControl_SE method, we developed a bias correction framework using gnomAD v3.1.2 AFs as proxies for the true AFs. We tested our framework using the Pan-UKB Diabetes datasets. The adjusted MAFs using the bias correction had less bias and a higher Lin’s CCC for both the AFR and EUR samples than the unadjusted method (**Figure 3, Table 2-3**). The average CCC across all variants improved from 0.9369 to 0.9877 for the PanUKBB AFR diabetes GWAS and from 0.9382 to 0.9943 for the EUR sample. While the overall bias is much lower, we do observe that the adjusted MAFs with the bias correction retain a similar trend to the original SE method with an increase in bias and variability as MAF increases.

**Figure 3.**
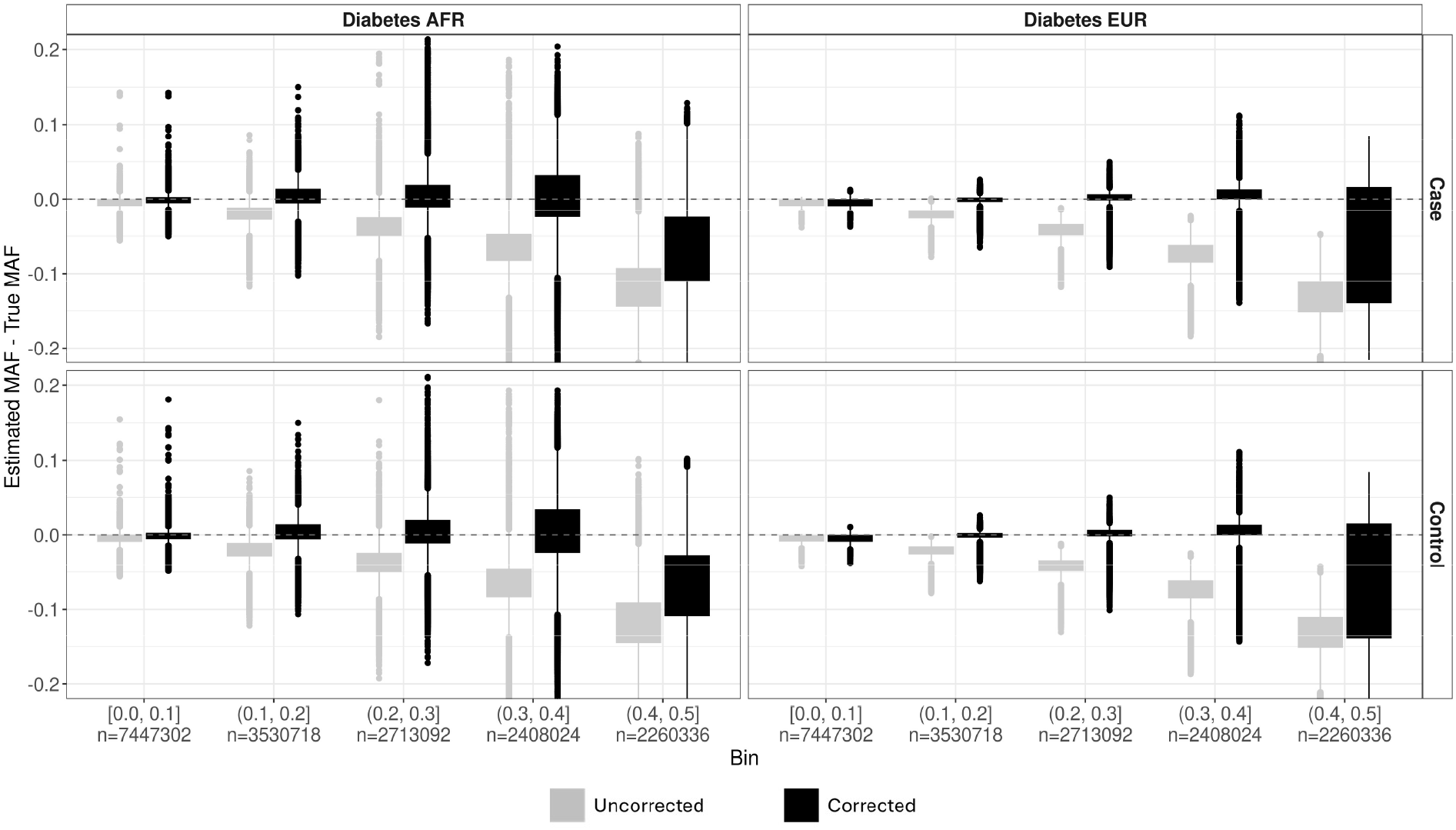
Correction mitigates bias in CaseControl_SE MAF estimates. We use our bias correction to adjust the case and control MAF estimates from CaseControl_SE for >9M genome-wide variants from the African and European Pan-UKB Diabetes datasets. To estimate the bias correction, we used >1.2M variants on chromosome 1 that were harmonized between Pan-UKBB and gnomAD v3.1.2. GnomAD non-Finnish European (NFE) were used as the proxy for true MAFs for the EUR sample (Right), and gnomAD African/African American (AFR/AFRAM) were used as the proxy for true MAFs for the AFR sample (Left). We see an improvement (i.e., less bias and greater Lin’s CCC) when using the bias correction framework (gray; AFR CCC = 0.9877, EUR CCC = 0.9943), compared to the uncorrected CaseControl_SE MAF estimates (black; AFR CCC = 0.9369, EUR CCC = 0.9382)

## 4 Discussion

Here, we present methods and an R package, *CCAFE*, to estimate case and control specific AFs from GWAS summary statistics. While our method using total AF outperforms using SE with less bias and variability in the estimates, the different methods allow flexibility based on what summary statistics are available. This is especially important given inconsistency in the summary statistics released in repositories such as the GWAS catalog (Buniello et al., 2019). Indeed, SE is often more available and easily derivable (e.g. given the effect estimate and either a p-value or test statistic) than total AF. To increase robust use of the SE method, we developed and provide an adjustment within the R package that greatly decreases the bias in the case and control AF estimates.

In both simulations and real data, we show that CaseControl_AF, provides highly accurate estimates of case and control AFs across a variety of sample sizes and in the presence of covariates. Conversely, while CaseControl_SE provides accurate AF estimates when no covariates are included in the original GWAS, this method is biased when the original GWAS includes covariates. Since CaseControl_AF is much less biased and more precise, we recommend using CaseControl_AF when total AFs are available. We hypothesize that the bias may arise from using the SE, which can be underestimated when covariates are included (Xing & Xing, 2010). Using the MAF results in more variability as well as results in a loss of information regarding the MAF and allele pairing, which can result in errors when harmonizing datasets. A key assumption of the method proposed by Yang *et al*. is that the allele being estimated is always the minor allele. To ensure the solution is bounded between zero and one, the SE based method assumes that the AF is being calculated on the minor allele. As such, only the MAF is output, and there is no connection between the reported alleles in the summary statistics and the estimated MAF. This loss of information regarding alleles can complicate secondary analyses, as inferring which allele is the minor or major allele may not be possible, especially when the values are close to 0.5. Conversely, our method using AF, CaseControl_AF, outputs the estimated case and control AF for which the total sample AF was reported, therefore retaining alternate and reference allele information.

As SE is more commonly available in GWAS summary data compared tototal AF, we developed a bias correction framework that provides more accurate case and control MAF estimates when the total sample AF is not available. The correction framework requires harmonization of the observed data with a publicly available data source such as gnomAD. However, we find that a subset of the data, such as chromosome one as we use here, can be used to estimate the bias correction relieving some of the additional computational and person-time burden and enabling bias correction for variants not in the public data. Future studies of the minimum number of overlapping variants between the observed and proxy data would be beneficial. While the bias correction improves the accuracy of the case and control AF estimates, variability and bias in higher MAF bins remains and is especially prominent when the sample size is small. Future development of a correction for this variability due to sample size would further improve the bias correction. Furthermore, to perform the bias adjustment, a publicly available database containing ancestrally matched proxy AFs is required, highlighting the need for large, public, ancestrally diverse databases. While gnomAD is an expansive resource for genomics, studies of diverse, specifically admixed, samples may not have an ancestrally matched gnomAD group to use as the proxy. For bias correction of the Pan-UK Biobank Diabetes GWAS in AFR individuals, we used gnomAD MAFs from the African/African American (AFR/AFRAM) group as proxies, and found that, while not a perfect ancestral substructure match, bias of the estimates was still greatly reduced. Summary data harmonization methods, such as *Summix2* (Stoneman et al., 2024), can adjust AFs to match genetic similarity between samples and could be used to harmonize the population structure between the proxy and GWAS data here. The use of this bias correction framework for ancestrally diverse or admixed samples requires additional investigation.

## 5 Conclusion

We have introduced methods and software to derive case and control AFs from GWAS summary data. The functions available in the *CCAFE* package provide researchers with user-friendly and open-source methods to enhance the re-use of publicly available genetic summary statistics.

## Data Availability

The datasets used here are available through Pan-UK Biobank Per-phenotype files (https://pan-dev.ukbb.broadinstitute.org/docs/per-phenotype-files/index.html) and the GWAS catalog for the prostate cancer GWAS (https://www.ebi.ac.uk/gwas/studies/GCST006085). The data from gnomAD is publicly available for download at (https://gnomad.broadinstitute.org/downloads). The package *CCAFE* is available for download here on GitHub: https://github.com/wolffha/CCAFE/ and through Bioconductor. Code used to generate simulated datasets and perform analyses is available here: https://github.com/wolffha/CaseControlAF_manuscript.

## Acknowledgements

This work was supported by the National Human Genome Research Institute [R35HG011293, R01HG011345, U01HG011715]. This work also used the computing resources at the Center for Computational Mathematics, University of Colorado Denver, including the Alderaan cluster, supported by the National Science Foundation award OAC-2019089.

