## Supplemental Material for "CCAFE: Estimating Case and Control Allele Frequencies from GWAS Summary Statistics"

**Supplement**

Full derivation:

Known:

$$AF_{total} = \frac{(N_{case}AF_{case} + N_{control}AF_{control})}{N_{total}} \tag{1}$$

$$OR = \frac{ad}{bc} \tag{2}$$

Where:

$a = 2N_{case} * AF_{case}$
$b = 2N_{case}(1 - AF_{case})$
$c = 2N_{control} * AF_{control}$
$d = 2N_{control}(1 - AF_{control})$

Find $AF_{control}$ and $AF_{case}$:

Substitute for $a, b, c, \& \ d$ into equation (2) and simplify:

$$OR = \frac{(2N_{case} * AF_{case})[2N_{control}(1 - AF_{control})]}{[2N_{case}(1 - AF_{case})](2N_{control} * AF_{control})} \tag{3}$$

$$OR = \frac{AF_{case}(1 - AF_{control})}{(1 - AF_{case})AF_{control}} \tag{4}$$

Substitute for $AF_{case}$ using equation (1) and distribute:

$$OR = \frac{(\frac{N_{total}}{N_{case}}AF_{total} - \frac{N_{control}}{N_{case}}AF_{control})(1 - AF_{control})}{[1 - \frac{N_{total}}{N_{case}}AF_{total} + \frac{N_{control}}{N_{case}}AF_{control}]AF_{control}} \tag{5}$$

$$OR = \frac{\frac{N_{total}}{N_{case}}AF_{total} - AF_{control}\frac{N_{control}}{N_{case}} - AF_{control}\left(\frac{N_{total}}{N_{case}}AF_{pop}\right) + AF_{control}^{2}(\frac{N_{control}}{N_{case}})}{AF_{control}[1 - \left(\frac{N_{total}}{N_{case}}AF_{total}\right)] + AF_{control}^{2}(\frac{N_{control}}{N_{case}})} \tag{6}$$

$$\begin{aligned} &AF_{control}^{2}\left(\frac{N_{control}}{N_{case}}\right)OR + AF_{control}\left[1 - \left(\frac{N_{total}}{N_{case}}AF_{total}\right)\right]OR \\ &\quad = AF_{control}^{2}\left(\frac{N_{control}}{N_{case}}\right) - AF_{control}\left(\frac{N_{control}}{N_{case}} + \frac{N_{total}}{N_{case}}AF_{total}\right) + \frac{N_{total}}{N_{case}}AF_{total} \end{aligned} \tag{7}$$

Arrange as quadratic equation:

$$AF_{control}^{2}\left[\frac{N_{control}}{N_{case}}(OR - 1)\right] + AF_{control}\left[OR\left(1 - \left(\frac{N_{total}}{N_{case}}AF_{total}\right)\right) + \frac{1}{N_{case}}(N_{control} + N_{total}AF_{total})\right] - \frac{N_{total}}{N_{case}}AF_{total} = 0 \tag{8}$$

Let:

$$\left[a = \frac{N_{control}}{N_{case}}(OR - 1)\right] \quad (9)$$

$$b = \left[OR\left(1 - \left(\frac{N_{total}}{N_{case}}AF_{total}\right)\right) + \frac{1}{N_{case}}(N_{control} + N_{total}AF_{total})\right] \quad (10)$$

$$c = -\frac{N_{total}}{N_{case}}AF_{total} \quad (11)$$

$$AF_{control}^2 a + AF_{control} b + c = 0 \quad (12)$$

Choose the root greater than 0 and less than 1 to be $AF_{control}$ where $x_1, x_2$ are the roots

$$x_{1,2} = \frac{-b \pm \sqrt{b^2 - 4ac}}{2a} \quad (13)$$

$$AF_{control} = \begin{cases} x_1, & if\ 0 \leq x_1 \leq 1 \\ x_2, & otherwise \end{cases}$$

Use the calculated $AF_{control}$ to solve for $AF_{case}$ where:

$$AF_{case} = \frac{N_{total}}{N_{case}}AF_{total} - \frac{N_{control}}{N_{case}}AF_{control} \quad (14)$$

Simulations show there is only one root [0,1].

Given the quadratic in equation (8) and the coefficients in (9 – 11), we sought to determine the possible solutions for the roots as shown in equation (13). Keeping $N_{total}$ constant at 10,000, we examined 144 different scenarios with varying AF, OR, $N_{case}$, and $N_{control}$ as shown in the table below (all combinations and results in Table S1).

| Parameter | Values |
|---|---|
| $N_{case}$ | 1000, 5000, 9000 |
| $N_{control}$ | 9000, 5000, 1000 |
| $N_{total}$ | 10000 |
| $AF_{total}$ | 0, 0.1, 0.25, 0.5, 0.75, 1 |
| $OR$ | 0.1, 0.4, 0.6, 0.8, 1.1, 1.2, 3, 5 |

We then used the values of the parameters to compute *a, b, c* as shown in equations (9-11), followed by calculation of the two roots using equation (13). Root 1 ($x_1$) and root 2 ($x_2$) were calculated as follows:

$$x_1 = \frac{-b - \sqrt{b^2 - 4ac}}{2a} \quad (15)$$

$$x_2 = \frac{-b + \sqrt{b^2 - 4ac}}{2a} \quad (16)$$

We observed the following results, indicating that for possible combinations of parameters, only one root ($x_2$) lies within [0,1]. Also note that when $a > 0$, $x_1 < 0$, and when $a < 0$, $x_1 > 1$.

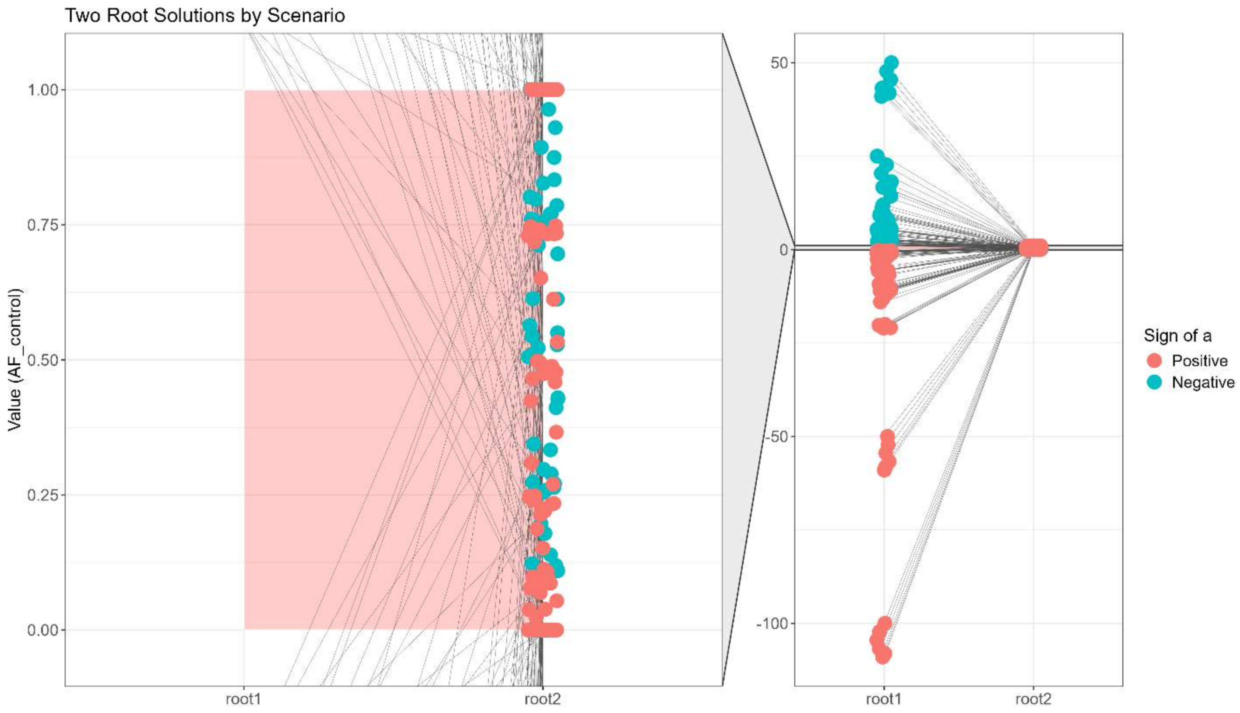


**Figure S1. Simulations show only one root solves for valid control AF value.** Simulations of 144 different scenarios with varying case and control sample size, OR, and AF were used to calculate the roots (two possible solutions for control AF). Root 2, calculated by equation (16) is always in the valid interval [0,1] (shown in the red rectangle), while root 1, calculated by equation (15) is always outside of the valid interval. We also note that when the coefficient *a* is positive, root 1 is lower than the valid interval (i.e. < 0) and when *a* is negative, root 1 is greater than the valid interval (i.e. > 1).

Case AF Derivation for Prostate Cancer GWAS

We selected a prostate cancer PGS based on a GWAS that published OR and control AF for each variant(Schumacher et al., 2018). Using the case and control sample sizes ($N_{case}, N_{control}$) we derive allele counts (AC) for the effect and non-effect alleles using $AC = AF * N$. We also know that the OR can be calculated using ACs as follows:

$$OR = \frac{AC_{effect,case} * AC_{non-effect,control}}{AC_{effect,control} * AC_{non-effect,case}} \tag{17}$$

We can then rearrange this equation to solve for the AC of the effect allele in cases, as needed for our reference data as follows:

$$AC_{effect,case} = \frac{OR * AC_{effect,control}}{1 - AC_{effect,control} + OR * AC_{effect,control}} \tag{18}$$

CaseControl_SE Bias Correction Framework

We observed systematic bias in the estimated case and control MAFs derived by CaseControl_SE in simulations with covariates and in real data. This bias was present in the estimated case, control, and total sample MAFs. To estimate this bias, we can use an estimate or proxy of the true MAF. We use gnomAD MAFs as this proxy. Because the bias trends with MAF bin, we fit regression models by MAF bin. For each MAF bin ([0, 0.1) [0.1, 0.2) [0.2, 0.3) [0.3, 0.4) [0.4, 0.5]) we fit a second order polynomial where the outcome is the estimated MAF from the SE method and the predictor is the gnomAD MAF (Figure S2).

We evaluated using smaller (5%) MAF bins and observed nearly identical least squares (LS) difference. We therefore chose 10% MAFs for computational efficiency. To obtain the best fitting model for the majority of the data, we identified and excluded outliers defined as observations with an absolute value of the studentized residual > 3. We then refit the models without outliers. This results in the regression estimates shown below.

$$\widehat{MAF}_{CaseControl\_SE} = \hat{a} * MAF_{gnomAD}^2 + \hat{b} * MAF_{gnomAD} + \hat{c} \quad (19)$$

The bias can then be estimated as:

$$\widehat{bias} = MAF_{gnomAD} - \widehat{MAF}_{CaseControl\_SE} \quad (20)$$

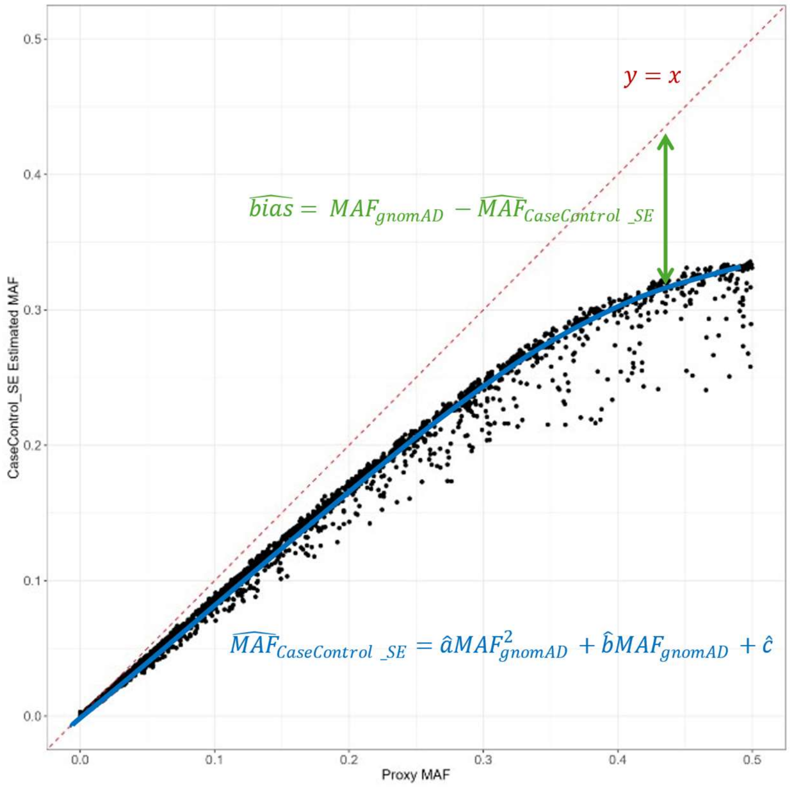


**Figure S2. Bias correction framework to adjust CaseControl_SE estimates using gnomAD as proxies.** The relationship between the proxy MAF (x-axis) and the estimated MAF (y-axis) is modeled through polynomial regressions (blue). This model is used to estimate the bias (green). The adjusted MAF estimate is estimated by adding the bias to the CaseControl_SE MAF output.

Since we observed a systematic bias where the estimated MAF was less than the true MAF, we estimate the bias by subtracting the predicted value from the gnomAD MAF. The adjusted MAF is calculated by adding the estimated bias to the estimated MAF output by CaseControl_SE. We found that using the same estimated bias for cases, controls, and total sample worked well.

$$MAF^{*}_{CaseControl\_SE} = MAF_{CaseControl\_SE} + \widehat{bias} \tag{21}$$

*Bias adjustment for variants not in proxy data*
To complete the bias adjustment for variants not present in the proxy data, the following steps are used:

1. Polynomial regression (second order) is fit using the proxy data ($MAF_{gnomAD}$) as the predictor and the CaseControl_SE MAF ($MAF_{CaseControl\_SE}$) as the outcome by gnomAD MAF bin, resulting in estimates for $\hat{a}, \hat{b}, \hat{c}$
2. For variants not in proxy dataset (i.e. CaseControl_SE MAF, but unknown gnomAD MAF), solve for the estimated gnomAD MAF ($\widehat{MAF}_{gnomAD}$) using $\hat{a}, \hat{b}, \hat{c}$

$$0 = \hat{a} * MAF^{2}_{gnomAD} + \hat{b} * MAF_{gnomAD} + (\hat{c} - MAF_{CaseControl\_SE}) \tag{22}$$

3. Calculate $\widehat{MAF}_{CaseControl\_SE}$

$$\widehat{MAF}_{CaseControl\_SE} = \hat{a} * \widehat{MAF}^{2}_{gnomAD} + \hat{b} * \widehat{MAF}_{gnomAD} + \hat{c} \tag{23}$$

4. Calculate $\widehat{bias}$:

$$\widehat{bias} = \widehat{MAF}_{gnomAD} - \widehat{MAF}_{CaseControl\_SE} \tag{24}$$

5. Calculate $MAF^{*}_{estimated}$ using equation (21)

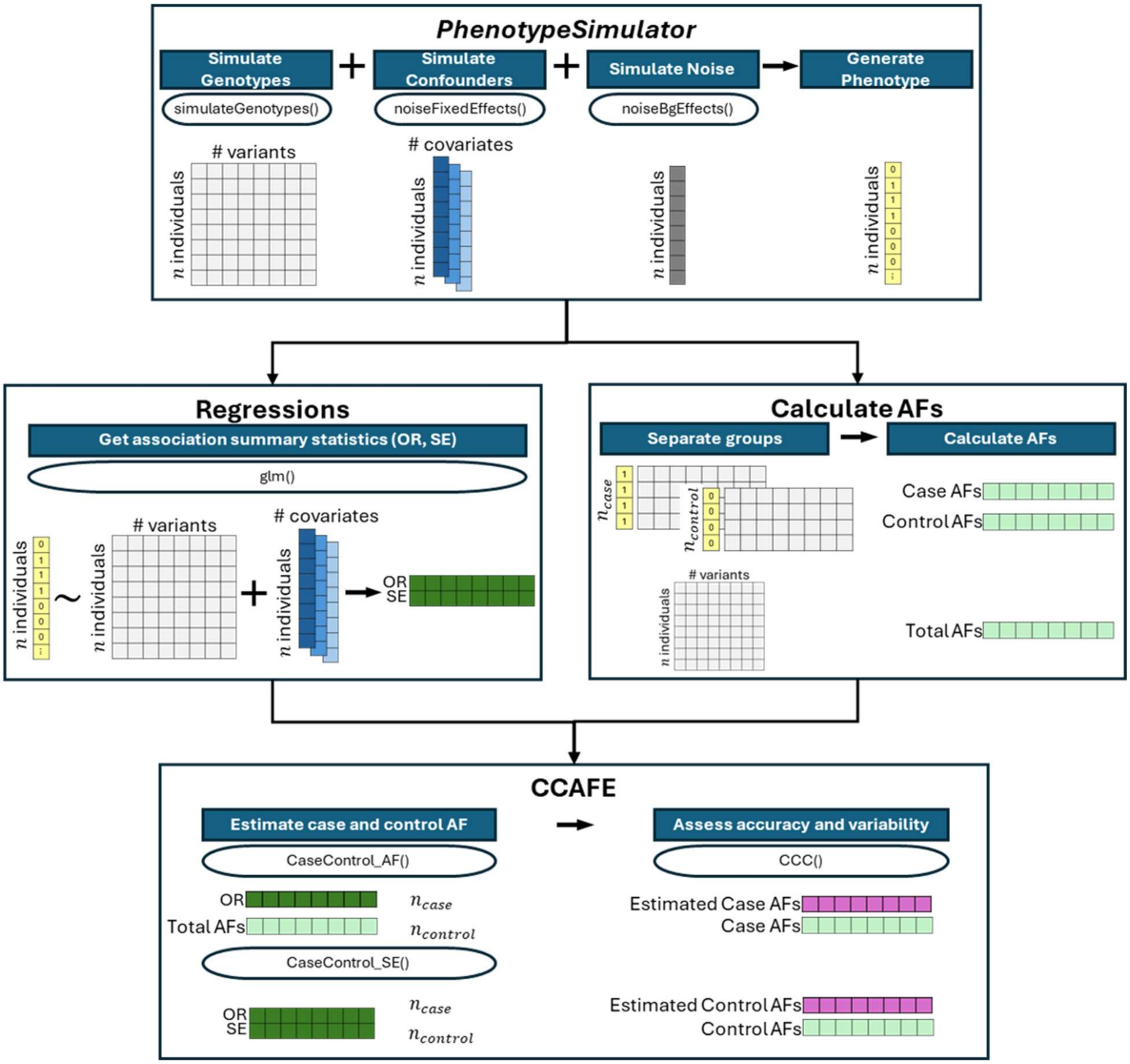


**Figure S3. Simulation framework to assess *CCAFE* accuracy and variability.** The *PhenotypeSimulator* R package was used to simulate genotypes, a binary phenotype, and covariates. Logistic regression was used on the simulated genotypes and phenotypes to obtain association summary statistics (OR, SE). The phenotype was also used to calculate AF for cases and controls. These simulations were used to test CaseControl_AF using the OR, total AF and case and control sample sizes as well as CaseControl_SE using the OR, SE, and case and control sample sizes. The output estimated case and control AFs were compared to the simulated case and control AFs using Lin's CCC.

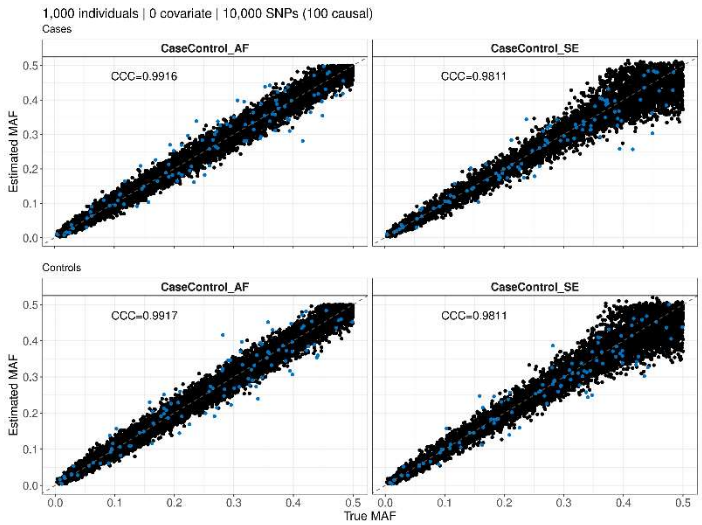


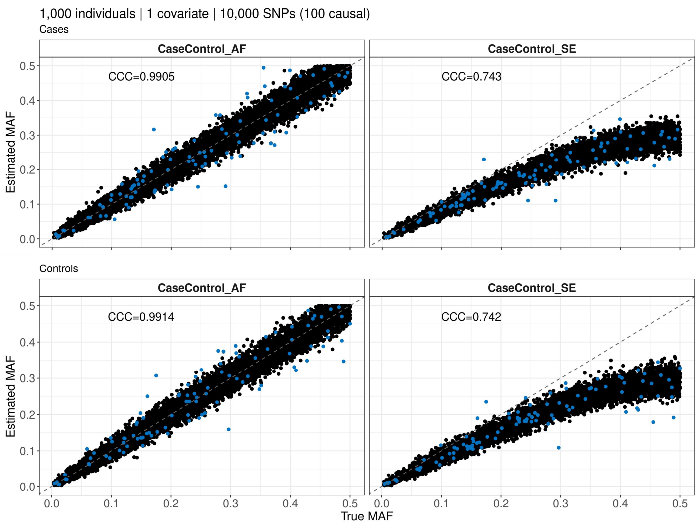


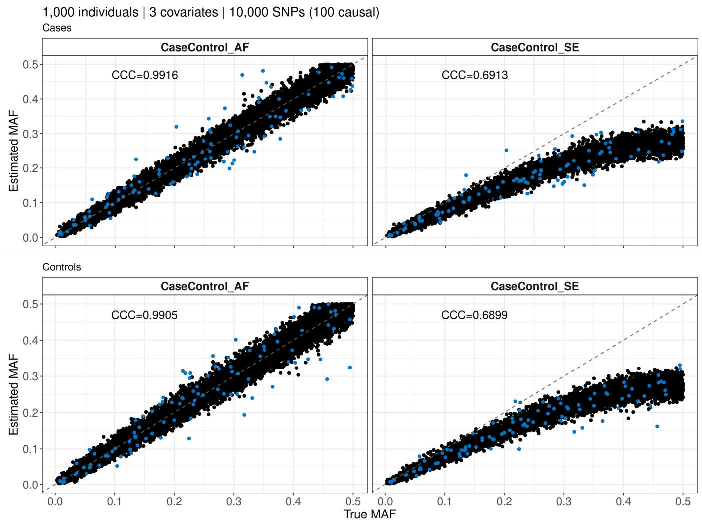


**Figure S4. CaseControl_SE produces biased results in simulation when covariates included.** Simulated genotypes and binary (case/control) phenotypes were generated using the PhenotypeSimulator R package for N=1,000 (500 cases and 500 controls). Genotypes for 10,000 SNPs, of which 100 were causal (blue), were generated for 1000 individuals. Logistic regression was used along with 0 (top left), 1 (top right), or 3 (bottom left) simulated covariates to generate summary statistics. The CCAFE R package was used to reconstruct the case and control AFs with total AF (left plots within each quadrant) or SE (right plots within each quadrant). Using the SE, bias was greater for higher MAFs and when more covariates were included. Using total AF was accurate across the simulation parameters evaluated. CCC values are reported in Table 1.

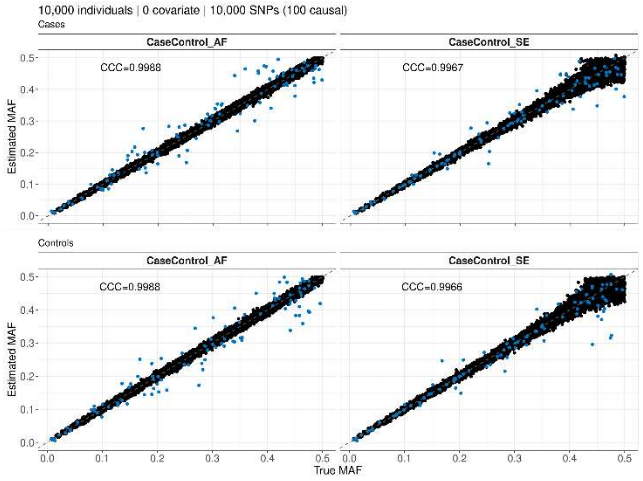


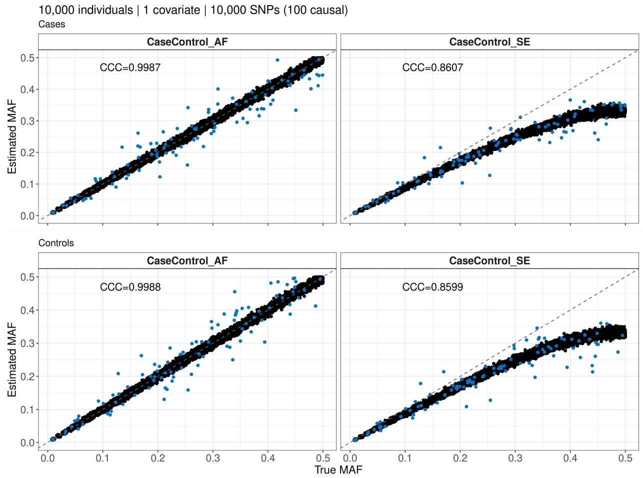


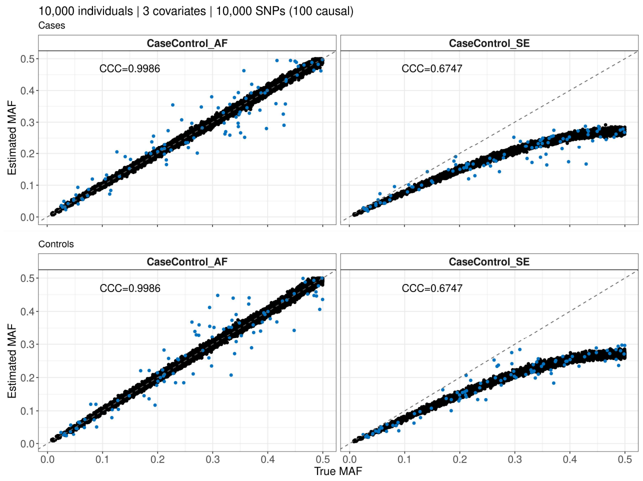


**Figure S5. CaseControl_SE produces biased results in simulation when covariates included.** Simulated genotypes and binary (case/control) phenotypes were generated using the PhenotypeSimulator R package for N=10,000 (5000 cases and 5000 controls). Genotypes for 10,000 SNPs, of which 100 were causal (blue), were generated for 1000 individuals. Logistic regression was used along with 0 (top left), 1 (top right), or 3 (bottom left) simulated covariates to generate summary statistics. The CCAFE R package was used to reconstruct the case and control AFs with total AF (left plots within each quadrant) or SE (right plots within each quadrant). Using the SE, bias was greater for higher MAFs and when more covariates were included. Using total AF was accurate across the simulation parameters evaluated. CCC values are reported in Table 1.

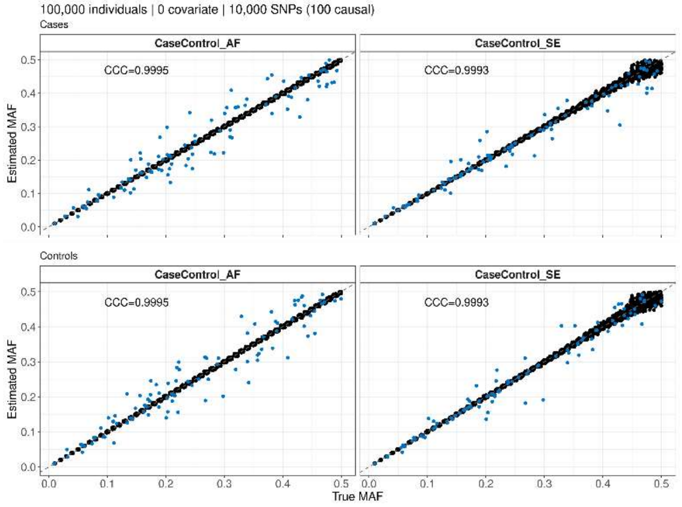


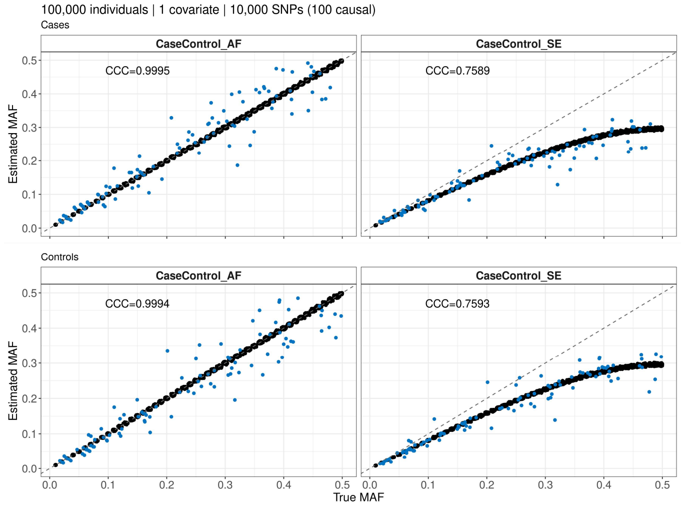


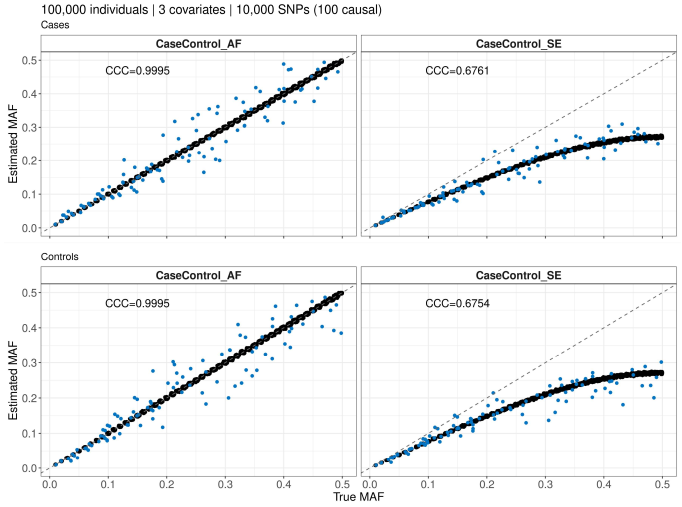


**Figure S6. CaseControl_SE produces biased results in simulation when covariates included.** Simulated genotypes and binary (case/control) phenotypes were generated using the PhenotypeSimulator R package for N=100,000 (50,000 cases and 50,000 controls). Genotypes for 10,000 SNPs, of which 100 were causal (blue), were generated for 1000 individuals. Logistic regression was used along with 0 (top left), 1 (top right), or 3 (bottom left) simulated covariates to generate summary statistics. The CCAFE R package was used to reconstruct the case and control AFs with total AF (left plots within each quadrant) or SE (right plots within each quadrant). Using the SE, bias was greater for higher MAFs and when more covariates were included. Using total AF was accurate across the simulation parameters evaluated. CCC values are reported in Table 1.

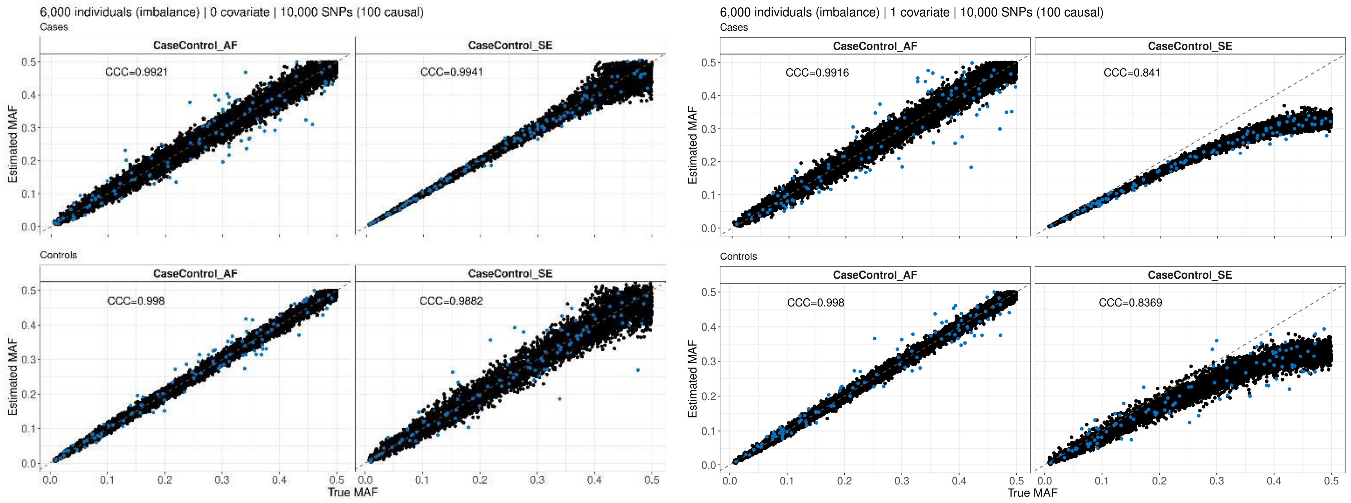


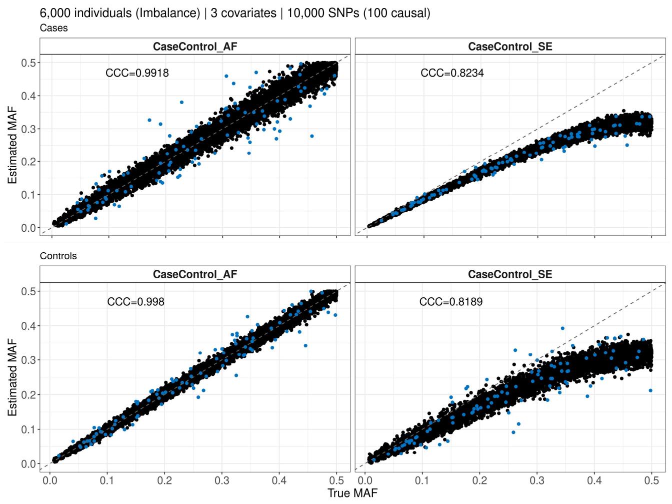


**Figure S7. CaseControl_SE produces biased results in simulation when covariates included and sample imbalance.** Simulated genotypes and binary (case/control) phenotypes were generated using the PhenotypeSimulator R package for N=6,000 (600 cases and 5,400 controls). Genotypes for 10,000 SNPs, of which 100 were causal (blue), were generated for 1000 individuals. Logistic regression was used along with 0 (top left), 1 (top right), or 3 (bottom left) simulated covariates to generate summary statistics. The CCAFE R package was used to reconstruct the case and control AFs with total AF (left plots within each quadrant) or SE (right plots within each quadrant). Using the SE, bias was greater for higher MAFs and when more covariates were included. Using total AF was accurate across the simulation parameters evaluated. CCC values are reported in Table 1.

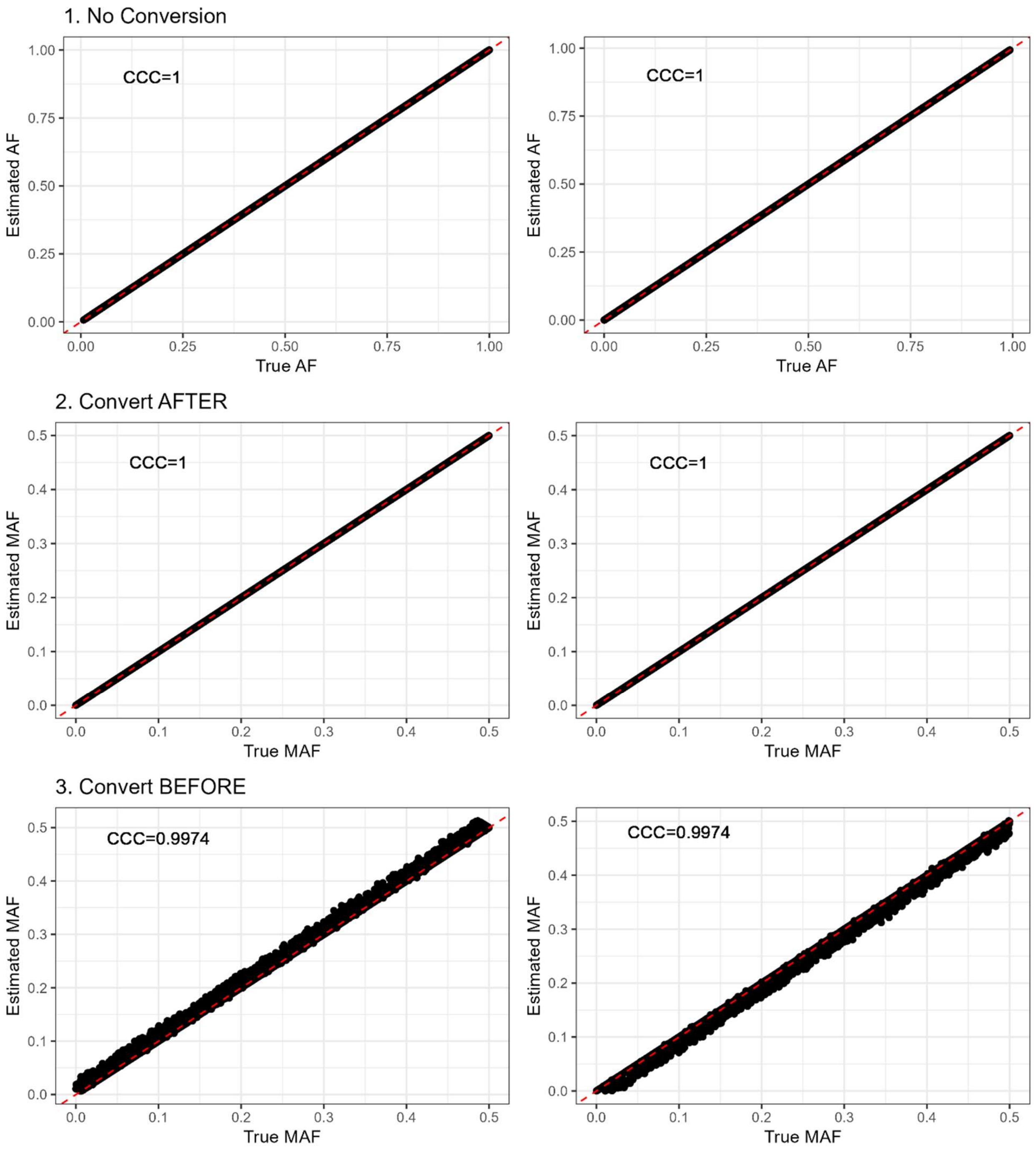


**Figure S8. Converting AF to MAF increases variability in case and control estimates using CaseControl_AF.** Using 2400 simulated variants three scenarios were tested to estimate case AF (Left) and control AF (Right) using CaseControl_AF: 1) using the total AF to estimate case and control AFs 2) using total AF to estimate case and control AFs then converting to minor AF (MAF) 3) converting total AF to MAF and estimating case and control MAFs. Converting first to MAF introduces variability. Lin's CCC is reported between the true and estimated AF.

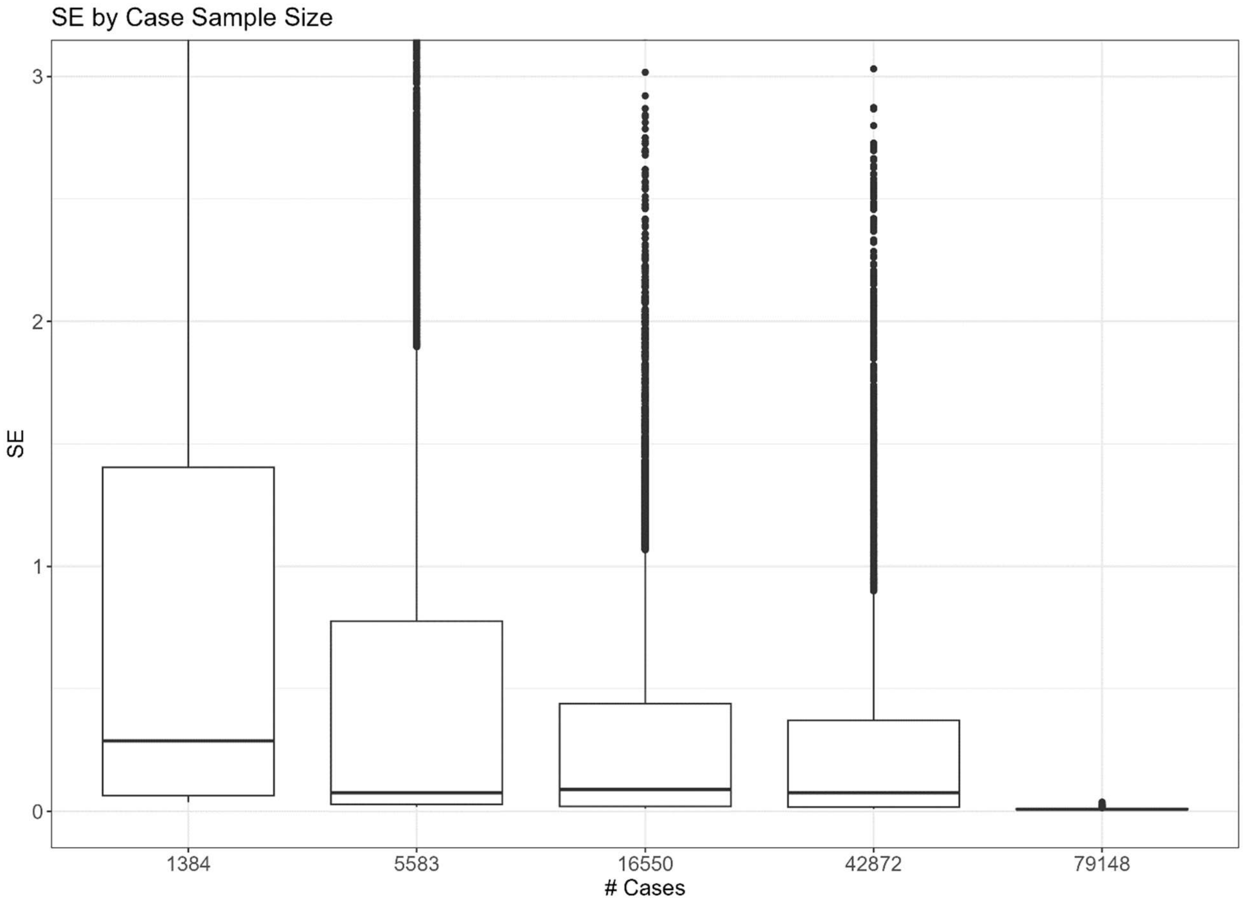


**Figure S9. Larger sample sizes have smaller SE and less variability in SE**. Here we examined the relationship between the case sample size and the standard error. We see studies with larger case sample sizes have a smaller SE including the minimum observed SE. Additionally, larger case sample sizes result in less variability in the SE across the variants. For each boxplot the center line shows the median SE and the upper and lower hinge represent the 75$^{th}$ and 25$^{th}$ percentile, respectively. The upper and lower whiskers are the largest and smallest values no more than 1.5* interquartile range (IQR) from the hinge.

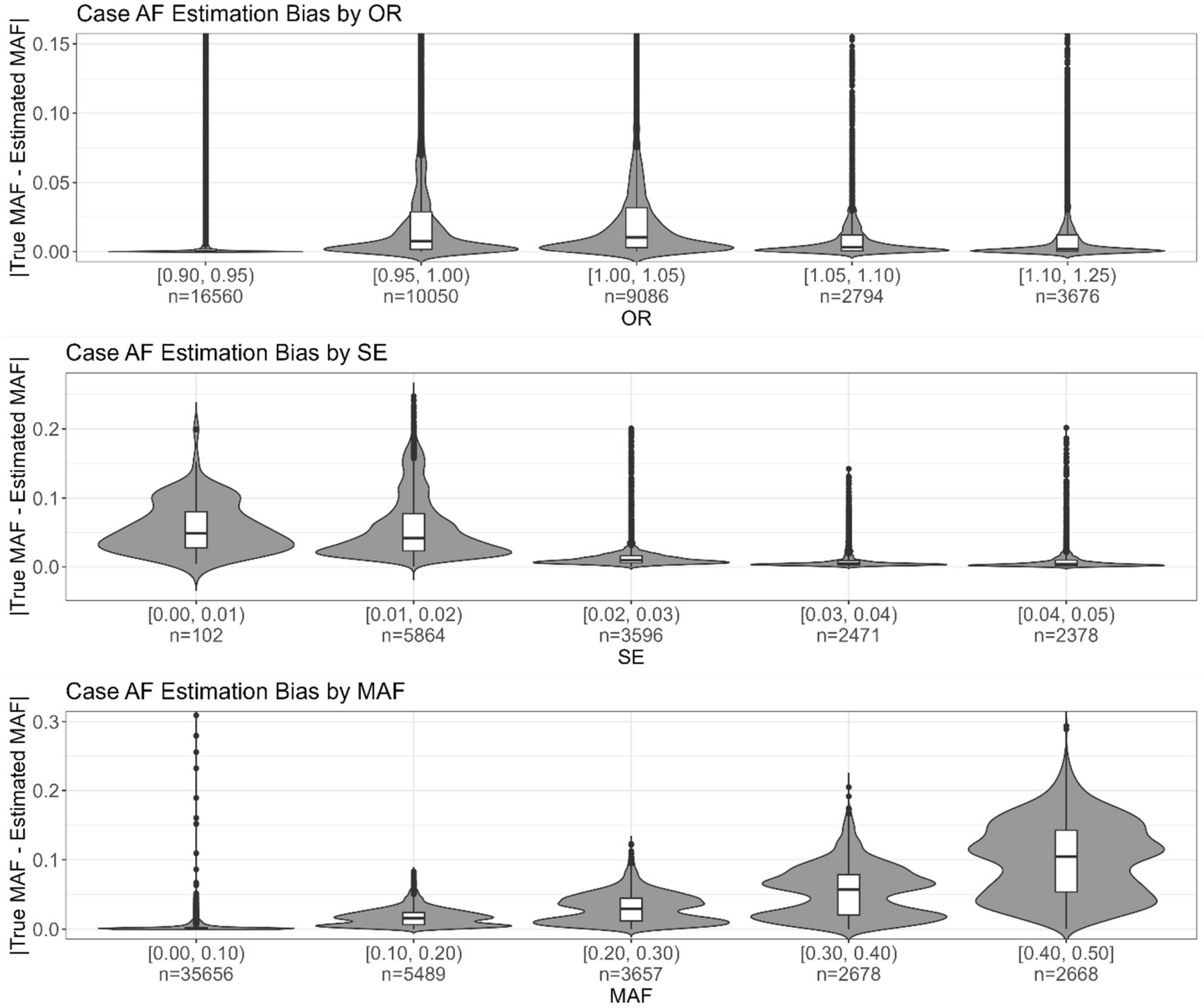


**Figure S10. CaseControl_SE has high bias at large MAF and small SE.** The difference between the true MAF and CaseControl_SE estimated MAF is compared across different bins for the OR (top), SE (middle), and MAF (bottom). Here the results for all datasets shown in Figure S7 are aggregated and plotted, with the number of SNPs in each bin shown on the x-axis. We see that bias is higher for smaller SE and larger MAF

| Table S1. Simulations show only one possible root (root2) falls within valid interval [0,1] for control AF solution | | | | | | | | | | |
|---|---|---|---|---|---|---|---|---|---|---|
| OR | AF | N_case | N_control | N_total | a | b | c | root1 | root2 | sim |
| 0.1 | 0 | 5000 | 5000 | 10000 | -0.9000 | 1.1000 | 0.0000 | 1.2222 | 0.0000 | 1 |
| 0.4 | 0 | 5000 | 5000 | 10000 | -0.6000 | 1.4000 | 0.0000 | 2.3333 | 0.0000 | 2 |
| 0.6 | 0 | 5000 | 5000 | 10000 | -0.4000 | 1.6000 | 0.0000 | 4.0000 | 0.0000 | 3 |
| 0.8 | 0 | 5000 | 5000 | 10000 | -0.2000 | 1.8000 | 0.0000 | 9.0000 | 0.0000 | 4 |
| 0.1 | 0 | 9000 | 1000 | 10000 | -0.1000 | 0.2111 | 0.0000 | 2.1111 | 0.0000 | 5 |
| 0.4 | 0 | 9000 | 1000 | 10000 | -0.0667 | 0.5111 | 0.0000 | 7.6667 | 0.0000 | 6 |
| 0.6 | 0 | 9000 | 1000 | 10000 | -0.0444 | 0.7111 | 0.0000 | 16.0000 | 0.0000 | 7 |
| 0.8 | 0 | 9000 | 1000 | 10000 | -0.0222 | 0.9111 | 0.0000 | 41.0000 | 0.0000 | 8 |
| 0.1 | 0 | 1000 | 9000 | 10000 | -8.1000 | 9.1000 | 0.0000 | 1.1235 | 0.0000 | 9 |
| 0.4 | 0 | 1000 | 9000 | 10000 | -5.4000 | 9.4000 | 0.0000 | 1.7407 | 0.0000 | 10 |
| 0.6 | 0 | 1000 | 9000 | 10000 | -3.6000 | 9.6000 | 0.0000 | 2.6667 | 0.0000 | 11 |
| 0.8 | 0 | 1000 | 9000 | 10000 | -1.8000 | 9.8000 | 0.0000 | 5.4444 | 0.0000 | 12 |
| 0.1 | 0.1 | 5000 | 5000 | 10000 | -0.9000 | 1.2800 | -0.2000 | 1.2435 | 0.1787 | 13 |
| 0.4 | 0.1 | 5000 | 5000 | 10000 | -0.6000 | 1.5200 | -0.2000 | 2.3941 | 0.1392 | 14 |
| 0.6 | 0.1 | 5000 | 5000 | 10000 | -0.4000 | 1.6800 | -0.2000 | 4.0774 | 0.1226 | 15 |
| 0.8 | 0.1 | 5000 | 5000 | 10000 | -0.2000 | 1.8400 | -0.2000 | 9.0900 | 0.1100 | 16 |
| 0.1 | 0.1 | 9000 | 1000 | 10000 | -0.1000 | 0.3111 | -0.1111 | 2.6995 | 0.4116 | 17 |
| 0.4 | 0.1 | 9000 | 1000 | 10000 | -0.0667 | 0.5778 | -0.1111 | 8.4699 | 0.1968 | 18 |
| 0.6 | 0.1 | 9000 | 1000 | 10000 | -0.0444 | 0.7556 | -0.1111 | 16.8516 | 0.1484 | 19 |
| 0.8 | 0.1 | 9000 | 1000 | 10000 | -0.0222 | 0.9333 | -0.1111 | 41.8806 | 0.1194 | 20 |
| 0.1 | 0.1 | 1000 | 9000 | 10000 | -8.1000 | 10.0000 | -1.0000 | 1.1248 | 0.1098 | 21 |
| 0.4 | 0.1 | 1000 | 9000 | 10000 | -5.4000 | 10.0000 | -1.0000 | 1.7458 | 0.1061 | 22 |
| 0.6 | 0.1 | 1000 | 9000 | 10000 | -3.6000 | 10.0000 | -1.0000 | 2.6739 | 0.1039 | 23 |
| 0.8 | 0.1 | 1000 | 9000 | 10000 | -1.8000 | 10.0000 | -1.0000 | 5.4537 | 0.1019 | 24 |
| 0.1 | 0.25 | 5000 | 5000 | 10000 | -0.9000 | 1.5500 | -0.5000 | 1.2923 | 0.4299 | 25 |
| 0.4 | 0.25 | 5000 | 5000 | 10000 | -0.6000 | 1.7000 | -0.5000 | 2.5000 | 0.3333 | 26 |
| 0.6 | 0.25 | 5000 | 5000 | 10000 | -0.4000 | 1.8000 | -0.5000 | 4.2026 | 0.2974 | 27 |
| 0.8 | 0.25 | 5000 | 5000 | 10000 | -0.2000 | 1.9000 | -0.5000 | 9.2291 | 0.2709 | 28 |
| 0.1 | 0.25 | 9000 | 1000 | 10000 | -0.1000 | 0.4611 | -0.2778 | 3.8986 | 0.7125 | 29 |
| 0.4 | 0.25 | 9000 | 1000 | 10000 | -0.0667 | 0.6778 | -0.2778 | 9.7388 | 0.4278 | 30 |
| 0.6 | 0.25 | 9000 | 1000 | 10000 | -0.0444 | 0.8222 | -0.2778 | 18.1558 | 0.3442 | 31 |
| 0.8 | 0.25 | 9000 | 1000 | 10000 | -0.0222 | 0.9667 | -0.2778 | 43.2107 | 0.2893 | 32 |
| 0.1 | 0.25 | 1000 | 9000 | 10000 | -8.1000 | 11.3500 | -2.5000 | 1.1275 | 0.2737 | 33 |
| 0.4 | 0.25 | 1000 | 9000 | 10000 | -5.4000 | 10.9000 | -2.5000 | 1.7547 | 0.2638 | 34 |
| 0.6 | 0.25 | 1000 | 9000 | 10000 | -3.6000 | 10.6000 | -2.5000 | 2.6859 | 0.2586 | 35 |
| 0.8 | 0.25 | 1000 | 9000 | 10000 | -1.8000 | 10.3000 | -2.5000 | 5.4682 | 0.2540 | 36 |
| 0.1 | 0.5 | 5000 | 5000 | 10000 | -0.9000 | 2.0000 | -1.0000 | 1.4625 | 0.7597 | 37 |
| 0.4 | 0.5 | 5000 | 5000 | 10000 | -0.6000 | 2.0000 | -1.0000 | 2.7208 | 0.6126 | 38 |
| 0.6 | 0.5 | 5000 | 5000 | 10000 | -0.4000 | 2.0000 | -1.0000 | 4.4365 | 0.5635 | 39 |
| 0.8 | 0.5 | 5000 | 5000 | 10000 | -0.2000 | 2.0000 | -1.0000 | 9.4721 | 0.5279 | 40 |
| 0.1 | 0.5 | 9000 | 1000 | 10000 | -0.1000 | 0.7111 | -0.5556 | 6.2176 | 0.8935 | 41 |

| | | | | | | | | | | |
|---|---|---|---|---|---|---|---|---|---|---|
| 0.4 | 0.5 | 9000 | 1000 | 10000 | -0.0667 | 0.8444 | -0.5556 | 11.9705 | 0.6962 | 42 |
| 0.6 | 0.5 | 9000 | 1000 | 10000 | -0.0444 | 0.9333 | -0.5556 | 20.3869 | 0.6131 | 43 |
| 0.8 | 0.5 | 9000 | 1000 | 10000 | -0.0222 | 1.0222 | -0.5556 | 45.4499 | 0.5501 | 44 |
| 0.1 | 0.5 | 1000 | 9000 | 10000 | -8.1000 | 13.6000 | -5.0000 | 1.1353 | 0.5437 | 45 |
| 0.4 | 0.5 | 1000 | 9000 | 10000 | -5.4000 | 12.4000 | -5.0000 | 1.7745 | 0.5218 | 46 |
| 0.6 | 0.5 | 1000 | 9000 | 10000 | -3.6000 | 11.6000 | -5.0000 | 2.7097 | 0.5126 | 47 |
| 0.8 | 0.5 | 1000 | 9000 | 10000 | -1.8000 | 10.8000 | -5.0000 | 5.4944 | 0.5056 | 48 |
| 0.1 | 0.75 | 5000 | 5000 | 10000 | -0.9000 | 2.4500 | -1.5000 | 1.7923 | 0.9299 | 49 |
| 0.4 | 0.75 | 5000 | 5000 | 10000 | -0.6000 | 2.3000 | -1.5000 | 3.0000 | 0.8333 | 50 |
| 0.6 | 0.75 | 5000 | 5000 | 10000 | -0.4000 | 2.2000 | -1.5000 | 4.7026 | 0.7974 | 51 |
| 0.8 | 0.75 | 5000 | 5000 | 10000 | -0.2000 | 2.1000 | -1.5000 | 9.7291 | 0.7709 | 52 |
| 0.1 | 0.75 | 9000 | 1000 | 10000 | -0.1000 | 0.9611 | -0.8333 | 8.6474 | 0.9637 | 53 |
| 0.4 | 0.75 | 9000 | 1000 | 10000 | -0.0667 | 1.0111 | -0.8333 | 14.2921 | 0.8746 | 54 |
| 0.6 | 0.75 | 9000 | 1000 | 10000 | -0.0444 | 1.0444 | -0.8333 | 22.6730 | 0.8270 | 55 |
| 0.8 | 0.75 | 9000 | 1000 | 10000 | -0.0222 | 1.0778 | -0.8333 | 47.7141 | 0.7859 | 56 |
| 0.1 | 0.75 | 1000 | 9000 | 10000 | -8.1000 | 15.8500 | -7.5000 | 1.1554 | 0.8014 | 57 |
| 0.4 | 0.75 | 1000 | 9000 | 10000 | -5.4000 | 13.9000 | -7.5000 | 1.8043 | 0.7698 | 58 |
| 0.6 | 0.75 | 1000 | 9000 | 10000 | -3.6000 | 12.6000 | -7.5000 | 2.7395 | 0.7605 | 59 |
| 0.8 | 0.75 | 1000 | 9000 | 10000 | -1.8000 | 11.3000 | -7.5000 | 5.5234 | 0.7544 | 60 |
| 0.1 | 1 | 5000 | 5000 | 10000 | -0.9000 | 2.9000 | -2.0000 | 2.2222 | 1.0000 | 61 |
| 0.4 | 1 | 5000 | 5000 | 10000 | -0.6000 | 2.6000 | -2.0000 | 3.3333 | 1.0000 | 62 |
| 0.6 | 1 | 5000 | 5000 | 10000 | -0.4000 | 2.4000 | -2.0000 | 5.0000 | 1.0000 | 63 |
| 0.8 | 1 | 5000 | 5000 | 10000 | -0.2000 | 2.2000 | -2.0000 | 10.0000 | 1.0000 | 64 |
| 0.1 | 1 | 9000 | 1000 | 10000 | -0.1000 | 1.2111 | -1.1111 | 11.1111 | 1.0000 | 65 |
| 0.4 | 1 | 9000 | 1000 | 10000 | -0.0667 | 1.1778 | -1.1111 | 16.6667 | 1.0000 | 66 |
| 0.6 | 1 | 9000 | 1000 | 10000 | -0.0444 | 1.1556 | -1.1111 | 25.0000 | 1.0000 | 67 |
| 0.8 | 1 | 9000 | 1000 | 10000 | -0.0222 | 1.1333 | -1.1111 | 50.0000 | 1.0000 | 68 |
| 0.1 | 1 | 1000 | 9000 | 10000 | -8.1000 | 18.1000 | -10.0000 | 1.2346 | 1.0000 | 69 |
| 0.4 | 1 | 1000 | 9000 | 10000 | -5.4000 | 15.4000 | -10.0000 | 1.8519 | 1.0000 | 70 |
| 0.6 | 1 | 1000 | 9000 | 10000 | -3.6000 | 13.6000 | -10.0000 | 2.7778 | 1.0000 | 71 |
| 0.8 | 1 | 1000 | 9000 | 10000 | -1.8000 | 11.8000 | -10.0000 | 5.5556 | 1.0000 | 72 |
| 1.1 | 0 | 5000 | 5000 | 10000 | 0.1000 | 2.1000 | 0.0000 | -21.0000 | 0.0000 | 73 |
| 1.2 | 0 | 5000 | 5000 | 10000 | 0.2000 | 2.2000 | 0.0000 | -11.0000 | 0.0000 | 74 |
| 3 | 0 | 5000 | 5000 | 10000 | 2.0000 | 4.0000 | 0.0000 | -2.0000 | 0.0000 | 75 |
| 5 | 0 | 5000 | 5000 | 10000 | 4.0000 | 6.0000 | 0.0000 | -1.5000 | 0.0000 | 76 |
| 1.1 | 0 | 9000 | 1000 | 10000 | 0.0111 | 1.2111 | 0.0000 | -109.0000 | 0.0000 | 77 |
| 1.2 | 0 | 9000 | 1000 | 10000 | 0.0222 | 1.3111 | 0.0000 | -59.0000 | 0.0000 | 78 |
| 3 | 0 | 9000 | 1000 | 10000 | 0.2222 | 3.1111 | 0.0000 | -14.0000 | 0.0000 | 79 |
| 5 | 0 | 9000 | 1000 | 10000 | 0.4444 | 5.1111 | 0.0000 | -11.5000 | 0.0000 | 80 |
| 1.1 | 0 | 1000 | 9000 | 10000 | 0.9000 | 10.1000 | 0.0000 | -11.2222 | 0.0000 | 81 |
| 1.2 | 0 | 1000 | 9000 | 10000 | 1.8000 | 10.2000 | 0.0000 | -5.6667 | 0.0000 | 82 |
| 3 | 0 | 1000 | 9000 | 10000 | 18.0000 | 12.0000 | 0.0000 | -0.6667 | 0.0000 | 83 |
| 5 | 0 | 1000 | 9000 | 10000 | 36.0000 | 14.0000 | 0.0000 | -0.3889 | 0.0000 | 84 |

| | | | | | | | | | | |
|---|---|---|---|---|---|---|---|---|---|---|
| 1.1 | 0.1 | 5000 | 5000 | 10000 | 0.1000 | 2.0800 | -0.2000 | -20.8957 | 0.0957 | 85 |
| 1.2 | 0.1 | 5000 | 5000 | 10000 | 0.2000 | 2.1600 | -0.2000 | -10.8918 | 0.0918 | 86 |
| 3 | 0.1 | 5000 | 5000 | 10000 | 2.0000 | 3.6000 | -0.2000 | -1.8539 | 0.0539 | 87 |
| 5 | 0.1 | 5000 | 5000 | 10000 | 4.0000 | 5.2000 | -0.2000 | -1.3374 | 0.0374 | 88 |
| 1.1 | 0.1 | 9000 | 1000 | 10000 | 0.0111 | 1.2000 | -0.1111 | -108.0925 | 0.0925 | 89 |
| 1.2 | 0.1 | 9000 | 1000 | 10000 | 0.0222 | 1.2889 | -0.1111 | -58.0861 | 0.0861 | 90 |
| 3 | 0.1 | 9000 | 1000 | 10000 | 0.2222 | 2.8889 | -0.1111 | -13.0383 | 0.0383 | 91 |
| 5 | 0.1 | 9000 | 1000 | 10000 | 0.4444 | 4.6667 | -0.1111 | -10.5238 | 0.0238 | 92 |
| 1.1 | 0.1 | 1000 | 9000 | 10000 | 0.9000 | 10.0000 | -1.0000 | -11.2102 | 0.0991 | 93 |
| 1.2 | 0.1 | 1000 | 9000 | 10000 | 1.8000 | 10.0000 | -1.0000 | -5.6538 | 0.0983 | 94 |
| 3 | 0.1 | 1000 | 9000 | 10000 | 18.0000 | 10.0000 | -1.0000 | -0.6421 | 0.0865 | 95 |
| 5 | 0.1 | 1000 | 9000 | 10000 | 36.0000 | 10.0000 | -1.0000 | -0.3558 | 0.0781 | 96 |
| 1.1 | 0.25 | 5000 | 5000 | 10000 | 0.1000 | 2.0500 | -0.5000 | -20.7411 | 0.2411 | 97 |
| 1.2 | 0.25 | 5000 | 5000 | 10000 | 0.2000 | 2.1000 | -0.5000 | -10.7329 | 0.2329 | 98 |
| 3 | 0.25 | 5000 | 5000 | 10000 | 2.0000 | 3.0000 | -0.5000 | -1.6514 | 0.1514 | 99 |
| 5 | 0.25 | 5000 | 5000 | 10000 | 4.0000 | 4.0000 | -0.5000 | -1.1124 | 0.1124 | 100 |
| 1.1 | 0.25 | 9000 | 1000 | 10000 | 0.0111 | 1.1833 | -0.2778 | -106.7342 | 0.2342 | 101 |
| 1.2 | 0.25 | 9000 | 1000 | 10000 | 0.0222 | 1.2556 | -0.2778 | -56.7204 | 0.2204 | 102 |
| 3 | 0.25 | 9000 | 1000 | 10000 | 0.2222 | 2.5556 | -0.2778 | -11.6077 | 0.1077 | 103 |
| 5 | 0.25 | 9000 | 1000 | 10000 | 0.4444 | 4.0000 | -0.2778 | -9.0689 | 0.0689 | 104 |
| 1.1 | 0.25 | 1000 | 9000 | 10000 | 0.9000 | 9.8500 | -2.5000 | -11.1926 | 0.2482 | 105 |
| 1.2 | 0.25 | 1000 | 9000 | 10000 | 1.8000 | 9.7000 | -2.5000 | -5.6353 | 0.2465 | 106 |
| 3 | 0.25 | 1000 | 9000 | 10000 | 18.0000 | 7.0000 | -2.5000 | -0.6148 | 0.2259 | 107 |
| 5 | 0.25 | 1000 | 9000 | 10000 | 36.0000 | 4.0000 | -2.5000 | -0.3249 | 0.2138 | 108 |
| 1.1 | 0.5 | 5000 | 5000 | 10000 | 0.1000 | 2.0000 | -1.0000 | -20.4881 | 0.4881 | 109 |
| 1.2 | 0.5 | 5000 | 5000 | 10000 | 0.2000 | 2.0000 | -1.0000 | -10.4772 | 0.4772 | 110 |
| 3 | 0.5 | 5000 | 5000 | 10000 | 2.0000 | 2.0000 | -1.0000 | -1.3660 | 0.3660 | 111 |
| 5 | 0.5 | 5000 | 5000 | 10000 | 4.0000 | 2.0000 | -1.0000 | -0.8090 | 0.3090 | 112 |
| 1.1 | 0.5 | 9000 | 1000 | 10000 | 0.0111 | 1.1556 | -0.5556 | -104.4786 | 0.4786 | 113 |
| 1.2 | 0.5 | 9000 | 1000 | 10000 | 0.0222 | 1.2000 | -0.5556 | -54.4591 | 0.4591 | 114 |
| 3 | 0.5 | 9000 | 1000 | 10000 | 0.2222 | 2.0000 | -0.5556 | -9.2697 | 0.2697 | 115 |
| 5 | 0.5 | 9000 | 1000 | 10000 | 0.4444 | 2.8889 | -0.5556 | -6.6869 | 0.1869 | 116 |
| 1.1 | 0.5 | 1000 | 9000 | 10000 | 0.9000 | 9.6000 | -5.0000 | -11.1643 | 0.4976 | 117 |
| 1.2 | 0.5 | 1000 | 9000 | 10000 | 1.8000 | 9.2000 | -5.0000 | -5.6066 | 0.4955 | 118 |
| 3 | 0.5 | 1000 | 9000 | 10000 | 18.0000 | 2.0000 | -5.0000 | -0.5855 | 0.4744 | 119 |
| 5 | 0.5 | 1000 | 9000 | 10000 | 36.0000 | -6.0000 | -5.0000 | -0.2985 | 0.4652 | 120 |
| 1.1 | 0.75 | 5000 | 5000 | 10000 | 0.1000 | 1.9500 | -1.5000 | -20.2411 | 0.7411 | 121 |
| 1.2 | 0.75 | 5000 | 5000 | 10000 | 0.2000 | 1.9000 | -1.5000 | -10.2329 | 0.7329 | 122 |
| 3 | 0.75 | 5000 | 5000 | 10000 | 2.0000 | 1.0000 | -1.5000 | -1.1514 | 0.6514 | 123 |
| 5 | 0.75 | 5000 | 5000 | 10000 | 4.0000 | 0.0000 | -1.5000 | -0.6124 | 0.6124 | 124 |
| 1.1 | 0.75 | 9000 | 1000 | 10000 | 0.0111 | 1.1278 | -0.8333 | -102.2336 | 0.7336 | 125 |
| 1.2 | 0.75 | 9000 | 1000 | 10000 | 0.0222 | 1.1444 | -0.8333 | -52.2181 | 0.7181 | 126 |
| 3 | 0.75 | 9000 | 1000 | 10000 | 0.2222 | 1.4444 | -0.8333 | -7.0332 | 0.5332 | 127 |

| | | | | | | | | | | |
|---|---|---|---|---|---|---|---|---|---|---|
| 5 | 0.75 | 9000 | 1000 | 10000 | 0.4444 | 1.7778 | -0.8333 | -4.4238 | 0.4238 | 128 |
| 1.1 | 0.75 | 1000 | 9000 | 10000 | 0.9000 | 9.3500 | -7.5000 | -11.1371 | 0.7482 | 129 |
| 1.2 | 0.75 | 1000 | 9000 | 10000 | 1.8000 | 8.7000 | -7.5000 | -5.5800 | 0.7467 | 130 |
| 3 | 0.75 | 1000 | 9000 | 10000 | 18.0000 | -3.0000 | -7.5000 | -0.5675 | 0.7342 | 131 |
| 5 | 0.75 | 1000 | 9000 | 10000 | 36.0000 | -16.0000 | -7.5000 | -0.2854 | 0.7299 | 132 |
| 1.1 | 1 | 5000 | 5000 | 10000 | 0.1000 | 1.9000 | -2.0000 | -20.0000 | 1.0000 | 133 |
| 1.2 | 1 | 5000 | 5000 | 10000 | 0.2000 | 1.8000 | -2.0000 | -10.0000 | 1.0000 | 134 |
| 3 | 1 | 5000 | 5000 | 10000 | 2.0000 | 0.0000 | -2.0000 | -1.0000 | 1.0000 | 135 |
| 5 | 1 | 5000 | 5000 | 10000 | 4.0000 | -2.0000 | -2.0000 | -0.5000 | 1.0000 | 136 |
| 1.1 | 1 | 9000 | 1000 | 10000 | 0.0111 | 1.1000 | -1.1111 | -100.0000 | 1.0000 | 137 |
| 1.2 | 1 | 9000 | 1000 | 10000 | 0.0222 | 1.0889 | -1.1111 | -50.0000 | 1.0000 | 138 |
| 3 | 1 | 9000 | 1000 | 10000 | 0.2222 | 0.8889 | -1.1111 | -5.0000 | 1.0000 | 139 |
| 5 | 1 | 9000 | 1000 | 10000 | 0.4444 | 0.6667 | -1.1111 | -2.5000 | 1.0000 | 140 |
| 1.1 | 1 | 1000 | 9000 | 10000 | 0.9000 | 9.1000 | -10.0000 | -11.1111 | 1.0000 | 141 |
| 1.2 | 1 | 1000 | 9000 | 10000 | 1.8000 | 8.2000 | -10.0000 | -5.5556 | 1.0000 | 142 |
| 3 | 1 | 1000 | 9000 | 10000 | 18.0000 | -8.0000 | -10.0000 | -0.5556 | 1.0000 | 143 |
| 5 | 1 | 1000 | 9000 | 10000 | 36.0000 | -26.0000 | -10.0000 | -0.2778 | 1.0000 | 144 |
